# Lysogeny destabilizes computationally simulated microbiomes

**DOI:** 10.1101/2023.10.23.563560

**Authors:** R. Tucker Gilman, Mark R. Muldoon, Spyridon Megremis, David L. Robertson, Nina Chanishvili, Nikolaos G. Papadopoulos

## Abstract

**Background:** The Anna Karenina Principle predicts that stability in host-associated microbiomes correlates with health in the host. Microbiomes are ecosystems, and classical ecological theory suggests that predators impact ecosystem stability. Phages can act as predators on bacterial species in microbiomes. However, our ability to extrapolate results from existing ecological theory to phages and microbiomes is limited because some phages can stage lysogenic infections, a process with no precise analog in classical ecology. In lysogenic infections, so-called “temperate” phages integrate into the cells of their hosts where they can remain dormant as prophages for many generations. Prophages can later be induced by environmental conditions, whereupon they lyse their host cells and phage progeny are released. It has been suggested that prophages can act as biological timebombs that destabilize microbial ecosystems, but formal theory to support this hypothesis is lacking.

**Results:** We studied the effects of temperate and virulent phages on diversity and stability in computationally simulated microbiomes. The presence of either phage type in a microbiome increased bacterial diversity. Bacterial populations were more likely to fluctuate over time when there were more temperate phages in the system. When microbiomes were disturbed from their steady states, both phage types affected return times, but in different ways. Bacterial species returned to their pre-disturbance densities more slowly when there were more temperate phage species, but cycles engendered by disturbances dampened more slowly when there were more virulent phage species.

**Conclusions:** Phages shape the diversity and stability of microbiomes, and temperate and virulent phages impact microbiomes in different ways. A clear understanding of the effects of phage life cycles on microbiome dynamics is needed to predict the role of microbiome composition in host health, and for applications including phage therapy and microbiome transplants. The results we present here provide a theoretical foundation for this body of work.

## Introduction

Understanding the factors that contribute to the stability of ecological communities is a fundamental goal of ecology [1–7]. Recently the topic has also drawn attention from practitioners in human health [8–10] and conservation [11, 12], largely due to increasing evidence for the Anna Karenina Principle (AKP). The AKP, proposed by Diamond [13] and first applied to microbial communities by Holmes and colleagues [9], rests on the well-supported observation that host-associated microbiomes are important contributors to the health of their hosts [14–17]. The AKP argues that each microbiome has a “healthy” state (i.e., composition) and that many different deviations from this state are associated with ill health [12]. This mirrors the opening statement of Tolstoy’s novel *Anna Karenina* that all happy families are alike, but every unhappy family is unhappy in its own way [18]. It is usually assumed that the healthy states of microbiomes are temporally stable, and that microbiomes depart from these states due to stochasticity or extrinsic perturbations [12]. Microbiomes with healthy states that are more strongly stable return to their healthy states more quickly after disturbances, and so spend less time in unhealthy states. Thus, stability in host-associated microbiomes may indicate or even help maintain health in the host [10–12].

The stability of ecological communities depends on how the species that make up those communities interact [3, 8, 19–21]. In simple model systems where all species are competitors, diversity is limited by competitive exclusion [6]. Systems with more competing species are more likely to show chaotic dynamics and return more slowly to their equilibria after disturbance [3], but the combined density of all species in these systems is less sensitive to disturbance than in systems with fewer competing species [3, 22]. If some species in a system are mutualists, then it is less likely that all species will persist in the stable state [8, 19], and if systems with mutualists are disturbed from their stable states they return more slowly than if all species were competitors [8, 23]. If a system includes predators, then predation can prevent prey species from reaching high densities. This reduces competition among prey, allowing more prey species to coexist and increasing prey diversity [24, 25]. Among systems with the same number of prey species, those that have more predators are more likely to show cyclic or chaotic dynamics and return more slowly to their steady states after disturbance [3]. Thus, the ecological roles of species in a microbiome may impact microbiome stability, and this in turn may impact host health.

Bacteriophages, often referred to simply as phages, are viruses that infect bacteria and thus act as predators on bacterial species in microbial ecosystems. There is growing interest in the use of phages to target disease-associated bacteria and influence the community composition of microbiomes for the management of human [26–30] and non-human [31–33] health. Extrapolating from models of other predator-prey systems, we might expect microbiomes with more phage species to have greater bacterial diversity and less stationary bacterial population dynamics than those with fewer phages. However, phages are unusual predators because they can have either lytic or lysogenic reproductive cycles [17, 34–36]. In the lytic cycle, the phage infects its bacterial host, the host cell is destroyed (i.e., lysed), and phage progeny are released. This is similar to attack by non-phage predators and approximates the assumptions of many classic predator-prey models [eg 2, 3, 6, 8, 19, 21]. In the lysogenic cycle, the phage infects its host but then integrates into the bacterial chromosome or persists in the host cell as a plasmid [35]. The phage (now called a prophage) can remain in this integrated state through many divisions of the bacterial host cell until some event or condition triggers induction [36]. Upon induction, the phage begins to replicate, the host cell is lysed, and phage progeny are released [35]. Both the probability that a phage enters the lysogenic cycle and the rate at which prophages are induced can depend on environmental factors including the densities of bacteria and phages [34, 37–39]. It has been suggested that lysogeny may alter the effect of phages on microbial ecosystems [32, 34, 35, 40–43], but little is known about how this mode of infection affects the diversity and dynamics of microbiomes [17, 30, 35, 44].

We studied the diversity and dynamic stability of computationally simulated microbiomes as part of the CURE project funded by the European Commission. Each simulated microbiome included a pool of bacterial species that competed for space or resources, and a set of phage species that could infect those bacterial species. We varied the numbers of virulent (i.e., obligately lytic) and temperate (i.e., potentially lysogenic) phage species among simulations. We simulated systems until they reached mature (i.e., stationary, cyclic, or chaotic) states. We asked how the diversity, and for nonstationary systems the temporal variability, of systems in the mature state depends on the numbers of virulent and temperate phage species in the system. Then, we disturbed the systems and asked how return times to the mature states depend on the numbers of virulent and temperate phage species. Our study represents the first attempt to model how phage life cycles affect the diversity and stability of rich bacterial communities.

## Methods

### Model

We simulated microbial systems in which bacterial species competed for space and resources following a Lotka-Volterra-like competition model (see supplementary information I.a). Each system had a species pool of 60 bacterial species. Bacteria of each species migrated into the system at constant rates, but some species were kept at low densities by competition. In many previous studies, researchers have drawn the strengths of competition between pairs of species independently from some distribution [eg 2, 3, 6, 8]. However, in nature, if a species competes strongly with two others, then it is likely that those two species also compete, and independent assignment of competition strengths does not capture such structured competition networks. To create ecologically plausible competition networks, we assigned four trait values to each bacterial species. These bacterial traits might represent the species’ optimal environmental conditions, degrees of dependence on particular resources, or any other attributes that cause more similar species to compete more strongly. We drew each trait value for each species independently from *N*(0,1). We set the strength of competition exerted by species *j* on species *i* to 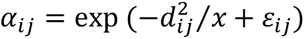, where 𝑑_𝑖𝑗_ is the Euclidean distance between species *i* and *j* in trait space, 𝑥 scales the strength of competition, and 𝜀_𝑖𝑗_ is drawn from *N*(0,0.01). The additive term 𝜀_𝑖𝑗_ ensured that the matrix of competition strengths was not strictly symmetrical. In nature competition is rarely strictly symmetrical [45], and models with strictly symmetrical competition can produce idiosyncratic behaviors [6, 46, 47]. We set 𝑥 to 1.2 in all simulations reported here, but qualitative results were similar for other values of 𝑥 that we investigated.

Each simulated system had a pool of up to 60 phage species that migrated into the system at constant rates. In the body of this paper we assumed that each phage species could infect only one bacterial species and that each bacterial species could be infected by only one phage species (i.e., phages are specialists). In the supplementary information we show that our results are qualitatively similar when phages can infect multiple bacterial species and bacterial species can be attacked by multiple phage species (i.e., phages are generalists; see supplementary information I.b, II.a). Each phage species in our system was either virulent or temperate. When a virulent phage infected a bacterium, the infection was lytic. The infecting phage was absorbed, the bacterial cell was lysed, and progeny phages were released. When a temperate phage infected a bacterium, the infection could be either lytic or lysogenic. In a lysogenic infection, the infecting phage was absorbed and the bacterial cell became a lysogen. In the body of this paper, we report results when lysogeny occurs with some fixed probability 𝑓 each time a temperate phage attacks a bacterium. In nature the probability of lysogeny can depend on the density of phages in the system [37, 38]. In the supplementary information we show that our qualitative results hold when the probability of lysogeny is density-dependent (see supplementary information I.c, II.b).

Once created, lysogens experienced density dependent population growth. Following Weitz [34], we assumed that lysogens compete on par with uninfected bacteria, but that their intrinsic population growth rates are reduced by a factor (1 − 𝑐), where 𝑐 can be interpreted as the cost experienced by bacteria that carry prophages. We assumed that lysogens cannot be attacked by the phage species they contain [48]. Lysogens excised phage DNA and returned to the uninfected state a low constant rate [34, 49]. Lysogens could be induced. When a lysogen was induced, the bacterial cell lysed and progeny of the infecting phage were released. In the body of this paper we assumed that induction occurs at a constant rate 𝑔, but in nature induction can depend on resource scarcity and thus on the density of competing bacteria in the system [34, 39, 50]. In the supplementary information we show that our qualitative results hold when the rate of induction is density-dependent (see supplementary information I.c, II.b).

The death of uninfected bacteria and lysogens from causes other than predation was captured by the competition-dependent population growth rates. Phages died at a constant rate that represents decay in the environment. This parameterization of predator and prey death is common in Lotka-Volterra-like models.

### Analysis

To understand how the properties of microbial systems depend on lysogeny, we studied 2 x 10^4^ systems with different numbers of virulent and temperate phage species. We simulated systems with different costs of lysogeny, lysogeny probabilities, and induction rates by controlling the parameters 𝑐, 𝑓, and 𝑔. For each simulation, we selected values for each of these parameters independently from 𝑐∼𝑈(0.1,0.9), 𝑓∼𝑈(0,1), and 𝑔 = 𝑒𝑥𝑝(𝑦) where 𝑦∼𝑈(−4.6,0). Under these parameter values, lysogeny can reduce bacterial growth rates by 10% to 90%, the probability of lysogenic attack for temperate phages can range from 0 to 1, and the induction rate of lysogens can range from one induction per generation to one induction per 100 generations. Thus, the patterns we observe in our full set of simulations emerge across a wide range of plausible systems.

We initialized our systems with all bacterial and phage population densities set to zero, and we allowed uninfected bacteria and phages to enter the systems by migration. We simulated each system in continuous time for 10^4^ time units. Under the parameter values we studied, this was sufficient for systems to reach mature states (i.e., stationary states, stable limit cycles, or persistent chaos). We estimated the time-average density for each uninfected bacterial and lysogen species over the last 10^3^ time units of each simulation, and we summed the time-average densities of the uninfected bacteria and lysogens of each species to obtain the mean total density for each bacterial species in the mature state of the system. We used the mean total densities of the bacterial species to compute the Shannon diversity of the bacterial community in the mature state. We counted the number of bacterial species with relative densities greater than 10^-4^ and called this the species richness of the mature system. We computed the coefficient of variation for each bacterial species, and we took the weighted mean coefficient of variation across all bacterial species to represent the temporal variability of the bacterial system in the mature state.

In addition to the characteristics of the mature states of our systems, we were interested in how quickly these systems return to their mature states after disturbances. To test this, we collected the final state of each system (i.e., the species densities after 10^4^ time units), and we perturbed the density of each uninfected bacterial species, phage species, and lysogen independently by multiplying each density by either *exp*(0.5) or *exp*(-0.5) with equal probability. We simulated the disturbed systems for 5 x 10^3^ time units and recorded two different return times. First, we recorded the return time to the mean population densities in the mature state. Specifically, we recorded the time from the disturbance until the midpoint of the first 200-time unit interval in which all time-averaged bacterial species densities greater than 10^-4^ were simultaneously within 1% of their pre-disturbance values. Second, we recorded the return time until population cycles triggered by the disturbance dampened. Specifically, we recorded the time from the disturbance until the midpoint of the first 200-time unit interval in which the weighted mean coefficient of variation was less than 10^-3^ or less than 1% greater than the weighted mean coefficient of variation in the mature state before the disturbance. We repeated this disturbance analysis three times for each simulated system.

To understand how lysogeny affects our response variables, we regressed each response variable on the number of virulent phage species in the system, the number of temperate phage species in the system, and the values of the system parameters 𝑐, 𝑓, and 𝑔. We included the interaction between the number of temperate phage species and the lysis rate (i.e., 1 − 𝑓) in each model. Including this interaction means that the regression coefficient associated with the number of temperate phage species describes the effect of phages that cause only lysogenic infections, which is a useful baseline for studying the effects of lysogeny.

Phage species richness and system parameters might affect the dynamical properties of bacterial systems directly, or they might affect bacterial diversity which might in turn affect the dynamical properties. We therefore conducted a second set of analyses in which we regressed each dynamical response variable on all the predictors in the first set of regressions, plus Shannon diversity, species richness, and (for return times) the weighted mean coefficient of variation over time for bacterial species in the mature system. This analysis tells us whether each predictor affects the dynamical properties of the system independent of any effects it might exert through its effect on bacterial diversity or dynamics in the mature state.

## Results

Systems with more phage species maintained more bacterial species and higher Shannon diversities of bacterial species (figure 1; table 1). Virulent phages had a greater effect on bacterial species richness (3.1% increase per phage species) than did temperate phages (1.9% increase per phage species). In the absence of phages, many bacterial species were excluded from systems by competition. Bacterial populations that were attacked by phages were maintained below their carrying capacities. This reduced competition and allowed more bacterial species to persist. Some bacteria that were infected by temperate phages survived as lysogens, so temperate phages had smaller effects on bacterial density and competition, and thus allowed fewer additional bacterial species to persist than did virulent phages. The relative effect of virulent and temperate phages on the Shannon diversity of bacterial species was more nuanced. When there were only a few virulent host-phage relationships in a system, the effect of temperate phages was greater than that of virulent phages (figure 1B, left side). When all or most of the bacterial species had virulent phage predators, the effect of virulent phages was greater than that of temperate phages (figure 1B, far right). The qualitative change in the relative effect of virulent and temperate phages corresponds to a change from bottom-up to top-down control of bacterial density in the systems (see supplementary information II.c).

**Figure 1.**
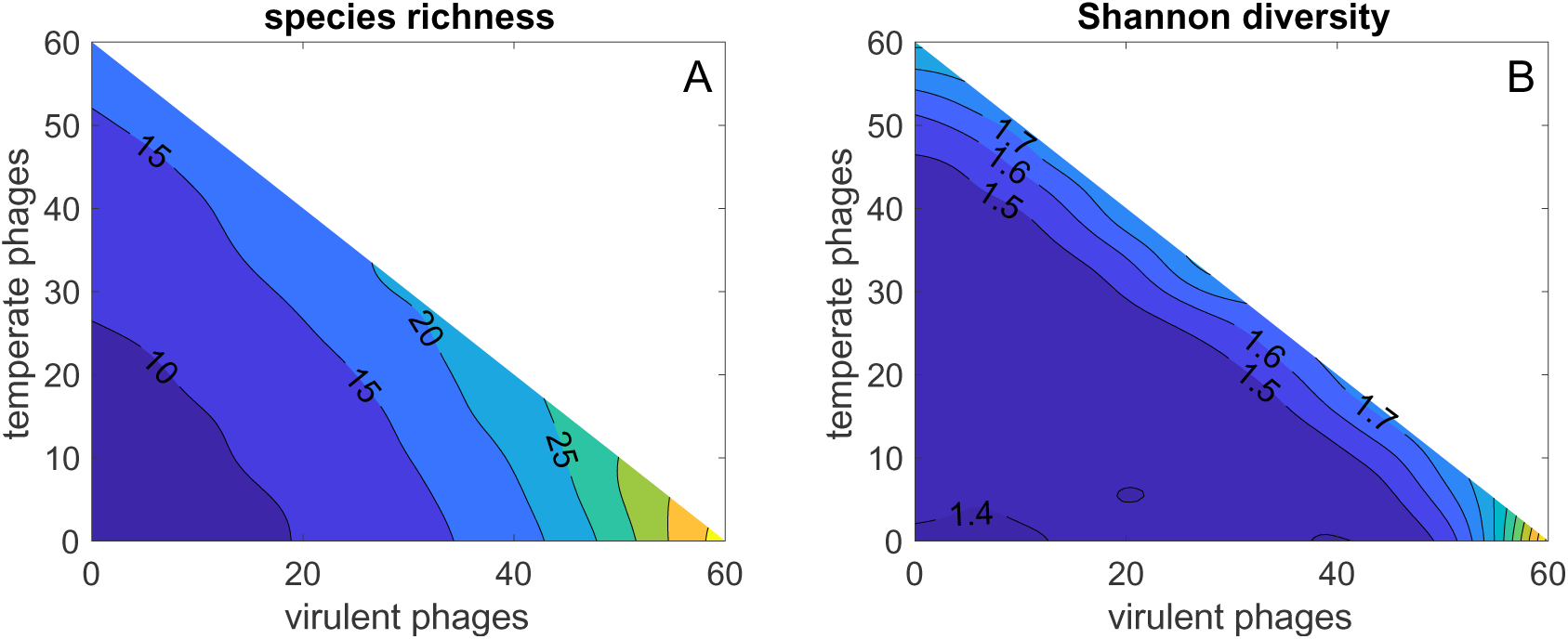
The mean species richness (A) and Shannon diversity (B) of bacterial communities in the mature state, as functions of the numbers of virulent and temperate phage species in the species pool. The upper triangles are not used because systems were capped at 60 phage species. Means are estimated over a Gaussian smoothing kernel (𝜎 = 3 phage species).

**Table 1.** The effects of the numbers of specialist virulent and temperate phage species in species pools and of the parameters that control lysogeny on the diversity and stability of bacterial communities. Effect sizes are reported as the change for each additional phage species. The effects on the probability that a system is nonstationary are reported as the change in log odds ratios. The top number in each cell indicates the effect of the predictor, and the bottom number indicates the effect of the predictor controlling for the diversity and variability of the bacterial community in the mature state. Nonsignificant effects are reported as ns.

| predictor | logged species richness | Shannon diversity | probability nonstationary | logged coefficient of variation (CoV) | logged return time to mean | logged return time to CoV |
| --- | --- | --- | --- | --- | --- | --- |
| number of phage species |  |  |  |  |  |  |
| virulent | 0.0308 | 0.0071 | 0.0255<br>-0.0678 | 0.0361<br>-0.0074 | -0.0013<br>-0.0134 | 0.0443<br>0.0318 |
| temperate | 0.0191 | 0.0080 | 0.0898<br>0.0620 | 0.0601<br>0.0257 | 0.0403<br>0.0287 | 0.0048<br>0.0050 |
| model parameters |  |  |  |  |  |  |
| cost of lysogeny (*c*) | 0.2942 | 0.1180 | 2.6044<br>1.9511 | 1.3532<br>1.0593 | 0.4468<br>0.2599 | 0.0492<br>ns |
| probability of lysogeny (*f*) | 0.0200 | ns | 5.3334<br>4.4083 | 1.4205<br>1.2474 | 0.1376<br>0.1544 | ns<br>-0.0809 |
| induction rate (*g*) | 0.7384 | 0.3072 | 9.3148<br>8.2580 | 1.9810<br>1.5590 | 1.3513<br>0.7418 | ns<br>-0.0550 |
| (1 - *f*) x number of temperate phage species | 0.0017 | -0.0017 | -0.0035<br>-0.0792 | ns<br>-0.0059 | -0.0295<br>-0.0240 | 0.0056<br>-0.0025 |
| emergent properties of systems |  |  |  |  |  |  |
| species richness |  |  | 0.1672 | 0.052 | 0.0269 | 0.0270 |
| Shannon diversity |  |  | -0.9631 | 0.242 | -0.3714 | 0.1523 |
| coefficient of variation (CoV) |  |  |  |  | 2.8790 | -3.1624 |

Among the systems we studied, 28.3% were non-stationary in the mature state. Temperate phages made bacterial dynamics more likely to be nonstationary (ΔLOR = 0.0898 per phage species) and increased the weighted mean coefficient of variance over time (6.2% increase per phage species) when bacterial dynamics were nonstationary (figure 2; table 1). Virulent phages also made bacterial dynamics more likely to be nonstationary (ΔLOR = 0.0255 per phage species) and increased temporal variability in systems that were nonstationary (3.7% increase per phage species), but this was mediated by changes in bacterial diversity. In mature systems with similar bacterial diversity, those with more temperate phages were more likely to be nonstationary (ΔLOR = 0.0620 per phage species) and were more variable when they were nonstationary (2.6% increase per phage species), but those with more virulent phages were less likely to be nonstationary (ΔLOR = -0.0678 per phage species) and were less variable if they were nonstationary (0.7% decrease per phage species).

**Figure 2.**
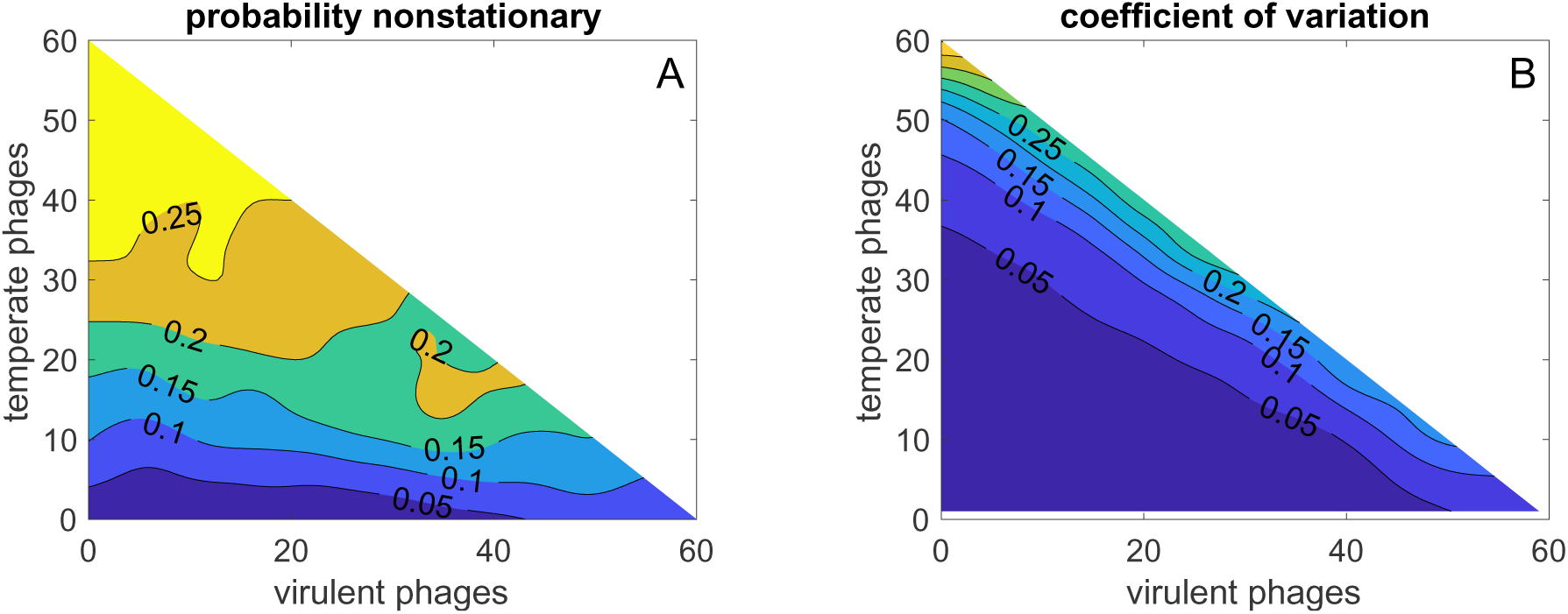
The probability that bacterial community dynamics were nonstationary (A) and the weighted mean coefficient of variation over time in nonstationary bacterial communities (B) in mature states as functions of the numbers of virulent and temperate phage species in the species pool. White areas of the lower triangle of (D) indicate that there were no nonstationary dynamics in the absence of temperate phages. Means are estimated over a Gaussian smoothing kernel (𝜎 = 3 phage species).

Bacterial species returned to their pre-disturbance densities more slowly in systems with more temperate phage species (4.1% increase in logged return time per species), but more quickly in systems with more virulent phages (0.1% decrease in logged return time per species; figure 3A; table 1). In systems with similar bacterial diversity, return times increased by 2.9% per temperate phage species and decreased by 1.3% per virulent phage species. Both temperate and virulent phages increased the time that it took for cycles started or amplified by disturbance to return to pre-disturbance amplitudes (figure 3B; table 1). However, unlike return times to pre-disturbance densities, the effect of virulent phages on return times to pre-disturbance amplitudes was greater than that of temperate phages (4.5% increase per virulent phage species vs 0.5% increase per temperate phage species). Systems with more virulent phages were less variable in the mature state than systems with more temperate phages (figure 2), but the greater effect of virulent than temperate phages on return times to pre-disturbance amplitudes is not because systems with more virulent phages were less variable and so had farther to return. Virulent phages increased return times to pre-disturbance amplitudes more than temperate phages even among simulations in which mature states were stationary (4.5% increase per virulent phage species vs 1.3% increase per temperate phage species; results not presented in table).

**Figure 3.**
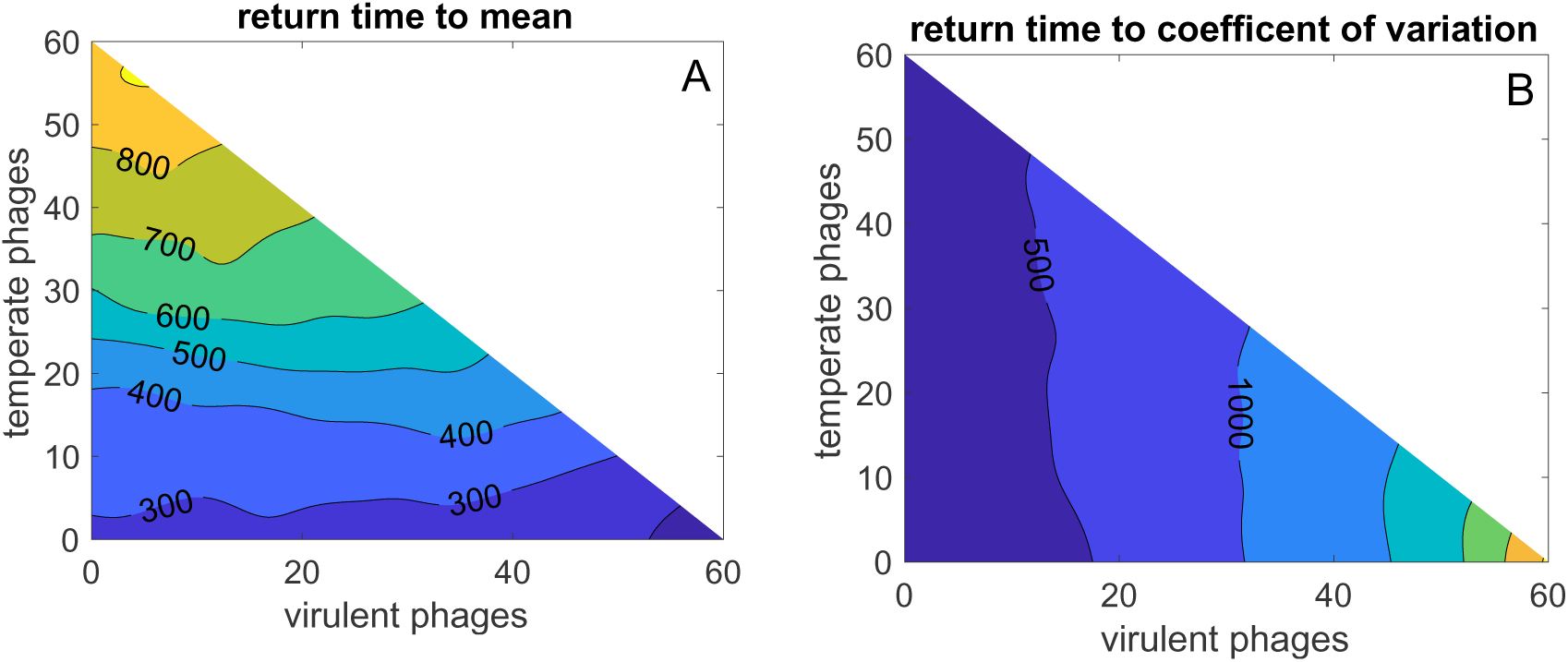
Expected return times of bacterial communities to pre-disturbance mean densities (A) and weighted mean coefficients of variation (B) after disturbance, as functions of the numbers of virulent and temperate phage species in the species pool. Means are estimated over a Gaussian smoothing kernel (𝜎 = 3 phage species).

The properties of lysogeny affected the diversity and stability of simulated microbiomes (table 1). When the growth rate of lysogens was slower (i.e., the cost of lysogeny for bacteria was higher), bacterial diversity in the mature state was higher, mature states were more variable over time, and return times were slower. As induction rates increased, bacterial diversity and variability in the mature states increased and return times to pre-disturbance densities increased but return times to pre-disturbance amplitudes decreased. As the probability of lysogeny decreased, temperate phages behaved more like virulent phages, and systems with more temperate phages behaved more like systems with more virulent phages.

## Discussion

Our results suggest that exposure to phages will typically increase the diversity of bacterial communities. The effect of virulent phages on bacterial diversity is likely to be greater than that of temperate phages. Exposure to phages will also increase the temporal variability of bacterial community composition. Communities with more phage species are more likely to have intrinsic cycles, and the amplitudes of those cycles are likely to be larger. This is partly because exposure to phages increases bacterial diversity, and systems with greater bacterial diversity are more temporally variable. Among systems with similar bacterial diversity, those with more temperate phage species are likely to be more variable over time, and those with more virulent phage species are likely to be less variable over time. If systems are disturbed from their mature states by extrinsic factors, those with more temperate phages are likely to return to their pre-disturbance densities more slowly. The effect of virulent phages on return times to pre-disturbance densities is smaller. Specialist virulent phages are likely to hasten return, but generalist virulent phages may slow return. Cycles engendered or amplified by extrinsic disturbances are likely to dampen more slowly when there are more phage species, and here the effects of temperate and virulent phages are reversed. Return times to pre-disturbance levels of temporal variability are likely to be slower when phages are virulent than when they are temperate.

The greater effects of temperate than virulent phages on the intrinsic temporal variability of bacterial systems is likely because lysogenic infections exert delayed effects on bacterial hosts. Temperate phages that successfully infect their hosts are removed from the active predator population, but hosts continue to compete and reproduce as lysogens, potentially for many generations, until some set of future environmental conditions triggers induction. May [6] argued more than 50 years ago that delays in Lotka-Volterra-like systems could lead to chaotic dynamics, although he did not apply this argument to temperate phages. More recently, it has been argued that temperate phages may serve as biological “time bombs” with the potential to suddenly and profoundly alter microbiome composition [51] and negatively impact the host health [41]. However, other researchers have argued that, at least in special cases, the induction of prophages can stabilize microbial communities [40]. Our model provides the first formal theoretical exploration of these conflicting hypotheses. The greater effect of virulent than temperate phages on return times of bacterial communities to pre-disturbance levels of temporal variability after extrinsic disturbances is likely because virulent phages are more efficient predators. Virulent infection can rapidly reduce bacterial populations and increase phage populations, causing systems to overshoot joint equilibria and leading to predator-prey cycles. When infections are temperate, infected bacteria are not destroyed immediately. Bacterial populations decline and phage populations increase more slowly, overshooting of equilibria is less severe, and cycles engendered or amplified by disturbances are less extreme and dampen more quickly.

The Anna Karenina Principle argues that microbiomes that are less variable or closer to their equilibria are more likely to be healthy for their hosts [11, 12]. In this context, one might wish to ask whether virulent or temperate phages maintain bacterial communities closer to their equilibria. Our study does not answer that question. We studied return times from single disturbances, but real systems are likely to experience repeated disturbances [52, 53]. These could be stochastic [53] or predictable over time (e.g., due to diurnal cycles or regulation by the host immune system) [52]. In the presence of repeated disturbances, it is not clear whether faster return to mean densities or faster dampening of cycles will keep systems closer to their equilibria, and the answer may depend on the disturbance regime. New theory that studies a range of biologically plausible disturbance regimes will be needed to answer this question. Furthermore, while the AKP posits that disturbed states of microbiomes are unhealthy, some disturbed states may be more unhealthy than others. New empirical work is needed to understand how different non-equilibrium states of microbiomes affect host health. The theory we present here will motivate and underpin that work.

Our study differs in an important way from some influential work on ecological stability. Many studies (e.g. [6, 8, 19, 21]; but see [2, 3]) have generated random ecological communities and then examined the special equilibria at which all species in those communities have positive densities. In these studies, “stability” means that all of the simulated species can coexist. This work has generally found that predator-prey interactions in a community increase the probability of coexistence [6, 19], consistent with our finding that systems with more phage predators maintain more bacterial species. Our approach differs from these earlier studies in that we allowed our randomly generated communities to relax to mature states before we studied them. We know that species present in the mature states of our systems can coexist or they would not be in the mature states. We studied instead whether the mature states are nonstationary or vary over time (i.e., due to chaos or persistent cycles), and how quickly the systems return to their mature states after perturbations. Because the AKP argues that temporal variability in host-associated microbiomes can indicate disease in the host, these measures of stability are useful for understanding how the composition of microbiomes might affect host health. In another model that studied relaxed states, Ives and Carpenter [3] found that among systems with the same number of prey species, those with more predators were more likely to show cyclic or chaotic dynamics and return more slowly to their mature states after disturbance. Our results extend this theory to virulent and temperate phages.

Our model makes many simplifying assumptions, but two are particularly worthy of note. First, we assumed that there was a maximum of one phage species that could lysogenize each bacterial species in the system. In practice, our model tracks each lysogen type separately, and if polylysogeny were possible in the model then the dynamical system we would need to simulate would be intractable for all but the smallest communities. Nonetheless, in nature, polylysogeny is not only possible but common [34, 36, 54]. New models are needed to test whether our results hold in systems with polylysogeny. Second, in our model, neither phages nor bacterial hosts evolve. In many macrobiological systems, ecological dynamics are faster than evolutionary dynamics, and models without evolution may be reasonable. In microbial systems, bacterial hosts may evolve resistance to phages, and phages may evolve to attack new bacterial species, on timescales that are ecologically relevant [54]. Host-phage coevolution can lead to the diversification of bacterial and phage species, with nested interaction structures in the newly evolved groups of strains [54–56]. Furthermore, because prophages rely on the survival and reproduction of their bacterial hosts to ensure their own survival and reproduction, temperate phages can evolve to benefit their lysogens by conferring increased growth rates under some conditions or by protecting their lysogens from attack by other phages [34, 54]. New work is needed to understand the dynamical behavior of microbiomes when bacteria and phages can coevolve.

## Conclusion

Phage therapy holds promise as a tool for managing host-associated microbial communities [26, 27, 31], and as bacterial resistance to antimicrobial drugs spreads globally [57, 58] interest in phage therapy is likely to grow [59–61]. Phage cocktails – engineered coalitions of phages designed to treat specific conditions – are particularly appealing for their potential to thwart the evolution of phage resistance in targeted bacterial strains [60]. Our results provide new evidence that coalitions of phages can profoundly impact the composition and dynamics of managed microbial systems. However, the impacts may be complex and difficult to predict, and the indiscriminate introduction of phages could push entire communities into states of dysbiosis. A deeper understanding of microbial dynamics and of phage potentials for lysogeny is needed to help researchers and practitioners deploy phage therapies safely and effectively. Our study offers a first step towards that goal.

## Supporting information

a.Codes and data for Gilman et al 2023.zip

## Supplementary information

### I. Supplementary methods

#### a. Model formalization

The dynamical system we studied can be written as

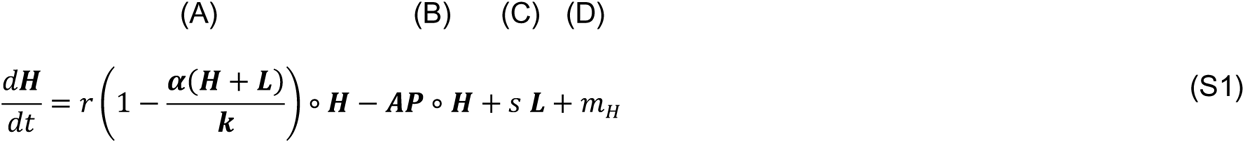

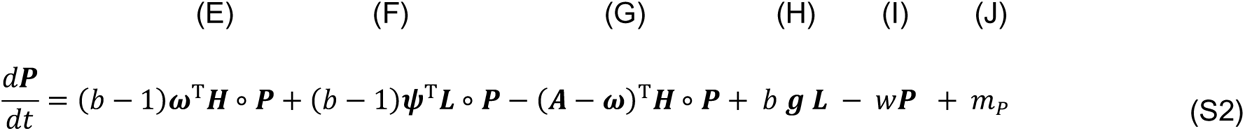

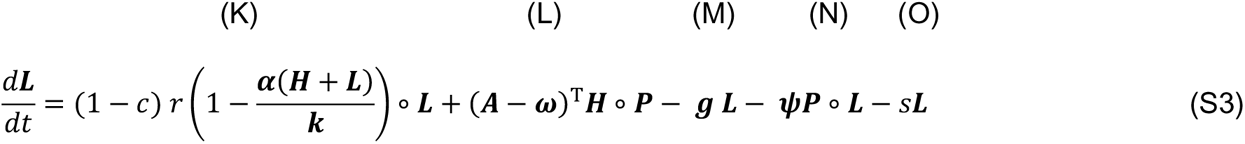

Here, ***H***, ***P***, and ***L*** are vectors that track the population densities of the uninfected bacterial, phage, and lysogen species, respectively, and the operator ∘ indicates the Hadamard product. ***H***, ***P***, and ***L*** are corresponding, such that entries ℎ_𝑖_ and 𝑙_𝑖_ track bacteria and lysogens of the same species, and 𝑝_𝑖_tracks the phage that produces lysogen 𝑖. In the body of the paper, we assumed that each phage species could infect only one bacterial species and that each bacterial species can be infected by only one phage species. In this supplementary material, we relax that assumption, so that phages may infect multiple bacterial species and bacterial species may have multiple phage predators (see section I.b). However, we continue to assume that each bacterial species can be lysogenized by at most one phage species, and therefore there is only one type of lysogen per bacterial species. In nature, polylysogeny is common[36], but models with large phage communities and polylysogeny are computationally intractable.

Equation S1 shows the instantaneous rates of population change for uninfected bacteria. Term (A) on the righthand side (RHS) of equation S1 captures competition-dependent population growth, where 𝑟 is the intrinsic population growth rate for uninfected bacteria, 𝒌 is the vector of carrying capacities for the bacterial species, and 𝜶 is the competition matrix in which entry 𝛼_𝑖𝑗_describes the competition exerted by bacterial species 𝑗 on bacterial species 𝑖. Term (B) captures the loss of uninfected bacteria due to infection by phages, where ***A*** is the predation matrix in which entry 𝑎_𝑖𝑗_ is the rate at which phages of species 𝑗 infect uninfected bacteria of species 𝑖. Term (C) captures the increase in the density of uninfected bacteria due to the spontaneous curing of lysogens at rate *s*. Term (D) captures the increase in the densities of uninfected bacteria species due to migration into the system at rate 𝑚_𝐻_. In principle, 𝑟, 𝑠, and 𝑚_𝐻_ could be vectors with different rates for each bacterial species, but for simplicity we assumed that all bacterial species share the same values for these parameters. We allowed carrying capacity to differ among bacterial species. In particular, 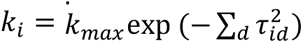 where 𝜏_𝑖𝑑_ is the trait value of bacterial species 𝑖 in dimension 𝑑 of trait space and 𝑘_𝑚𝑎𝑥_ scales the total bacterial carrying capacity of the system. Because bacterial trait values in our model are normally distributed around zero, this assignment of carrying capacities means that bacteria are more likely to have trait values that allow them to access common resource types than rare resource types.

Equation S2 shows the instantaneous rates of population change for phage species. Term (E) on the RHS of equation S2 captures the increase in phage densities due to the lytic infection of previously uninfected bacteria. Here, *b* is the burst size (ie, the number of phages released when a bacterial cell is lysed), 𝑏 − 1 accounts for the infecting phage that is absorbed in the infection, and 𝝎 is the matrix of lytic infection rates in which entry 𝜔_𝑖𝑗_ is the lytic infection rate of phage species 𝑗 on bacterial species 𝑖. For virulent phages and for temperate phages infecting bacteria that they cannot lysogenize, 𝜔_𝑖𝑗_ = 𝑎_𝑖𝑗_. If phage species 𝑗 is temperate and thus can lysogenize bacterial species 𝑗, then 𝜔_𝑗𝑗_ = 𝑎_𝑗𝑗_(1 − 𝑓_𝑗_) where 𝑓_𝑗_is the probability of lysogeny for phage species *j*. In the body of the paper we assumed that 𝑓_𝑗_ = 𝑓 for all phage species and that 𝑓 is constant in any given system. In this supplementary material, we show that results are qualitatively similar when the probability of lysogeny is density dependent and varies among phage species (see section I.c). Term (F) on the RHS of equation S2 captures the increase in phage densities due to the infection of lysogens. We assumed that lysogens cannot be attacked by the species of phage that they carry as a prophage, but that they can be infected by other phages that can infect the corresponding bacteria. Then, 𝝍 is the matrix of infection rates on lysogens, where 𝜓_𝑖𝑗_ = 𝑎_𝑖𝑗_ for 𝑖 ≠ 𝑗 and 𝜓_𝑖𝑖_ = 0. Term (G) captures the decrease in phage density when phages infect previously uninfected bacteria to form lysogens. The matrix (𝑨 − 𝝎) is the matrix of infection rates less the matrix of lytic infection rates, and is thus the matrix of lysogenic infection rates. Term (H) captures the increase in phage densities due to the induction of lysogens. In the vector 𝒈, entry 𝑔_𝑖_ is the induction rate of lysogen 𝑖. In the body of the paper we assumed that 𝑔_𝑖_ = 𝑔 for all phage species and that 𝑔 is constant in any given system. In this supplementary material, we show that results are qualitatively similar when the induction rate is density dependent and varies among phage species (see section I.c). Term (I) captures the decay of phages in the environment at rate 𝑤. Term (J) captures the migration of phages into the system at rate 𝑚_𝑃_. In principle, 𝑏, 𝑤, and 𝑚_𝑃_ could be vectors with different values for each phage species, but for simplicity we assumed that all phage species share the same values for these parameters.

Equation S3 shows the instantaneous rates of population change for lysogens of each bacterial species. Term (K) on the RHS of equation S3 captures competition-dependent population growth and is analogous to term (A) in equation S1, where 𝑐 measures the reduction in the intrinsic population growth rate for lysogens compared to that of uninfected bacteria. Term (L) captures the creation of lysogens when uninfected bacteria are infected by temperate viruses. Term (M) captures the loss of lysogens due to induction. Term (N) captures the loss of lysogens due to infection by other phages. Term (O) captures the loss of lysogens due to spontaneous curing at rate 𝑠. In principle, 𝑐 could be a vector that describes different effects of lysogeny on population growth for different bacterial species, but for simplicity we assumed lysogeny affects population growth in the same way for all bacterial species.

Our goal was to study how the numbers of temperate and virulent phage species and the properties of lysogeny affect the diversity and stability of bacterial communities. To focus on these effects, we fixed the model parameters that are not directly involved in lysogeny at plausible values, and we studied ranges of values for the parameters that are directly involved in lysogeny (table S1). We did not systematically investigate the effects of our fixed parameters on bacterial communities, but in spot checks of other parameter combinations, results were qualitatively similar to those we report here.

#### b. Modelling generalist phages

In the body of this paper, we studied systems in which each phage species infects only one bacterial species. In nature, many phages can infect multiple bacterial species [56]. Therefore, we conducted additional simulations to understand whether our results hold for generalist phages. In these simulations, we assumed that each phage has one bacterial species to which it is best adapted and that it infects at rate 𝑎_𝑚𝑎𝑥_. In particular, we assumed that phage 𝑗 is best adapted to bacterial species 𝑗, which is the bacterial species it can lysogenize if it is temperate. With probability *z*, each phage is also able to infect each other bacterial species in the system. If phage 𝑗 can infect bacterial species 𝑖, it does so at rate 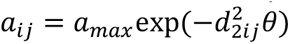, where 𝑑_2𝑖𝑗_ is the Euclidean distance between bacterial species 𝑖 and 𝑗 in the first two dimensions of the bacterial trait space. Thus, the ecological traits of bacterial species affect the rates at which they are infected by phages, but not in the same what that they affect competition. The parameter 𝜃 controls how strongly bacterial ecology affects phage infection attack rates. Under this formalization, we can control the parameter *z* to control whether phages in the model are specialists (*z* = 0) or generalists (*z* > 0). We conducted 2 x 10^4^ simulations with 𝑧∼𝑈(0,0.2) and 𝜃 = 0.1 and analyzed the behavior of the systems as in the body of the paper.

#### c. Modelling density dependent lysogeny and induction

In the body of this paper, we assumed that lysogeny occurs with the same fixed probability and induction occurs at the same constant rate for all phage and lysogen species. In nature, probabilities of lysogeny [37, 38] and induction rates [34, 39] can be density dependent. We conducted additional simulations to understand whether our results hold when this is true.

The relationships between phage density, bacterial density, lysogeny probability, and induction rate are not well characterized and may differ among systems. Therefore, we simulated systems with ranges of plausible relationships. In nature, the probability of lysogeny when a phage infects a bacterium can increase with the density of phages in the system [37, 38]. We set the probability of lysogeny when temperate phage 𝑖 attacks a bacterial species it can lysogenize to

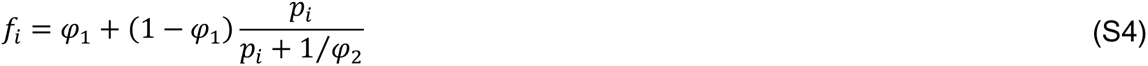

Here, 𝜑_1_ is the probability of lysogeny when the phage is rare and 𝜑_2_ controls how quickly the probability of lysogeny increases with phage density. In nature, the induction of lysogens can be triggered by strong resource competition [34, 39, 50]. To simulate this, we set the vector of induction rates for lysogen species to

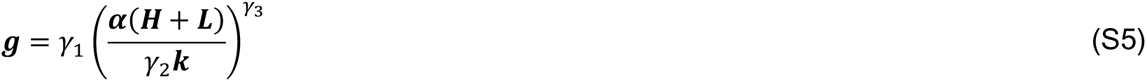

Here, 𝛾_1_ is a reference rate of induction, 𝛾_2_ controls the strength of competition at which the reference rate is reached, and 𝛾_3_ controls how rapidly the induction rate is increasing when the reference rate is reached. This formulation is sufficiently flexible to capture a wide range of biologically plausible relationships between resource competition and induction rate (figure S1).

We conducted 2 x 10^4^ simulations with 𝜑_1_∼𝑈(0,1), 𝜑_2_∼𝑈(0,2)^2^, 𝛾_1_∼𝐸𝑥𝑝(𝑈(−4.6,0)), 𝛾_2_∼𝑈(0.2,0.8), and 𝛾_3_∼𝑈(0,5). We analyzed the behavior of these systems as in the body of the paper, with one exception. In the body of the paper, we included the interaction between the number of temperate phage species and the lysis rate (1 − 𝑓) in our regression. This ensured that the regression coefficients associated with the number temperate phage species in the system were estimated for species that were always lysogenic, which is a useful baseline. When lysogeny rates are density dependent, the lysis rate varies among species and over time, and we cannot compute the lysis rate in the mature state from 𝜑_1_ and 𝜑_2_. Therefore, we observed the mean lysis rate across all potentially lysogenic infections in the mature state of the system, and we used this observed lysis rate rather than (1 − 𝑓) in our regression models.

The set of simulations described here is not an attempt to accurately capture any particular system in nature. However, if the qualitative results from our simple model hold in these systems, then this suggests that the results hold for many plausible forms of density dependent lysogeny and induction.

### II. Supplementary results

#### a. Results when phages are generalists

Figure S2 illustrates how the diversity (A, B), dynamics (C, D), and return times after disturbance (E, F) for bacterial communities depend on the numbers of virulent and temperate phage species in the species pool when phages are generalists (see also table S2). As when phages were specialists, bacterial species richness increased more rapidly with the number of virulent than temperate phages (figure S2A). Shannon diversity also increased with the number of phage species present (figure S2B). The relative rate of increase with the two phage types depended on the total number of phage species in the pool (see supplementary results II.c). Dynamics in the mature state were more likely to be nonstationary when there were more temperate phage species but not when there were more virulent phage species (figure S2C). If dynamics were nonstationary, variability over time was greater when there were more phage species, and the effect of temperate phage species was larger than that of virulent phage species (figure S2D). Return times after disturbance increased with the number of phage species in the species pool. Temperate phages had a greater effect on return times to pre-disturbance densities (figure S2E) and virulent phages had a greater effect on return times to pre-disturbance coefficients of variation (figure S2F).

The patterns we uncovered for generalist phages are qualitatively similar to those we reported for specialist phages in the body of the paper, but the differences between the phage types are less pronounced. In our model, each phage species was capable of lysogenizing only one bacterial species. Therefore, when temperate were generalists, their infections of many bacterial hosts were lytic, and thus their effects were more similar to those of virulent phages. In nature, some phages are capable of lysogenizing more than one bacterial host species [43, 44]. New theory is needed to understand how such generalist lysogeny affects bacterial community dynamics.

#### b. Results when the probability of lysogeny and the induction rate are density dependent

Figure S3 illustrates how the diversity (A, B), dynamics (C, D), and return times after disturbance (E, F) for bacterial communities depend on the numbers of virulent and temperate phage species when probabilities of lysogeny and induction rates are density dependent (see also table S3). Bacterial species richness increased more rapidly with the number of virulent than temperate phage species (figure S3A). Shannon diversity increased with the number of phage species (figure S3B), but the relative rate of increase with the two phage types depended on the total number of phage species in the pool (see supplementary results II.c). Dynamics in the mature state were more likely to be nonstationary when there were more temperate phage species, but less likely to be nonstationary when there were more virulent phage species in the species pool (figure S3C). If dynamics were nonstationary, temporal variability was greater when there were more phage species, and the effect of temperate phage species was larger than that of virulent phage species (figure S3D). Return times to pre-disturbance densities increased with the number of temperate phage species but decreased with the number of virulent phage species in the species pool (figure S3E). Return times to pre-disturbance coefficients of variation increased with the numbers of both phage types, but the effect of virulent phages was greater (figure S3F). The patterns reported here are qualitatively similar to those reported in the body of the paper for specialist phages when the probability of lysogeny and the induction rate were not density dependent.

#### **c.** The effects of phages on the Shannon diversity of bacterial communities

The relationship between the numbers of virulent and temperate phage species and the Shannon diversity of bacterial communities was more complicated than the other relationships we studied and requires additional explanation.

We begin by considering the simulations in which phage species were generalists (ie, the results presented in supplementary results II.b rather than in the body of the paper). Across all simulations with generalist phages, the Shannon diversities of bacterial communities in their mature states were bimodal (figure S4A). The residuals of the regression model we fit to Shannon diversity show two distinct clusters (figure S4B). This indicates that our regression model poorly explains Shannon diversity, underestimating the diversity of some systems and overestimating the diversity of others. Figure S4C shows the number of phage species in the species pool when bacterial communities had different Shannon diversities. High Shannon diversities occurred only when there were sufficiently large numbers of phage species in the pool. The redness of markers in figure S4C indicates the number of virulent phages in the species pool. Systems represented by redder markers had more virulent phage species. The predominance of red markers at the far right of figure S4C indicates that the highest Shannon diversities occurred when there were many virulent phage species available to the system.

The modes in Shannon diversity shown in figure S4A correspond to systems under bottom-up and top-down control. In the absence of phages, bacterial populations are limited by competition. Most systems are dominated by a small number of competitively dominant bacterial species, and other bacterial species are excluded from the system. Thus, bacterial diversity in these systems is low. If a virulent phage that can infect one of the dominant bacterial species enters the system, then the density of that bacterial species is reduced. In most cases, another bacterial species that is not subject to infection arises and becomes dominant, and the densities of most bacterial species remain limited by competition. The species richness of the bacterial community increases because one formerly excluded bacterial species is now common, but the Shannon diversity, which depends on both the richness and the evenness of bacterial species, changes very little. However, when *every* potentially dominant bacterial species can be infected by at least one virulent phage species, the control of the bacterial population densities switches from bottom-up (ie, controlled by competition) to top-down (ie, controlled by predation). This produces a step change in the evenness, and thus in the Shannon diversity, of the bacterial community. In figure S4A, the first mode corresponds to systems with few phage species, bottom-up control, and low bacterial diversity. The second mode corresponds to systems with many phage species, top-down control, and high bacterial diversity.

Because our initial regression models poorly described the Shannon diversities of bacterial communities when we studied all simulations together, we divided our results into two subsets – those with Shannon diversities less than 2.4, and those with Shannon diversities greater than 2.4. A Shannon diversity of 2.4 is the approximate nadir between the two modes in figure S4A. We fit new regression models to each subset of the data. The residuals of the fitted models no longer showed distinct clusters (figure S4D,E), suggesting that these new models are reasonable descriptions of the Shannon diversity within each subset of the data.

Table S3 shows the results of the new regressions. When Shannon diversities were less than 2.4 (ie, when bacterial populations experienced bottom-up control), the effect of temperate phage species on the Shannon diversity of the bacterial community was greater than the effect of virulent phage species. This is because temperate phages reduced the densities of their bacterial hosts and allowed competitors of those host species to increase, but they did not reduce the densities of the species on which they specialized so greatly that those hosts became rare. However, when phage species were common enough that all bacterial hosts had predators, then no bacterial species could reach carrying capacity and exclude the others, and there was a step change in Shannon diversity. At this point, control of the system switched to top-down, and the effect of virulent phages became stronger than that of temperate phages because virulent phages were more effective at limiting the densities of their bacterial hosts and so maintaining even bacterial species densities across the community.

The same mechanisms that operate when phage species are generalists also operate when they are specialists. However, the effects were less obvious because, when phage species were specialists, systems switched to top-down control only when almost all bacterial species had specialist phage predators, and this was the most extreme case we studied. Nonetheless, the patterns that arose when phages were generalists held when they were specialists. Regardless of whether the probabilities of lysogeny and induction rates were density dependent, Shannon diversities in the mature states of systems were bimodal (figures S5A and S6A) and there were distinct clusters of residuals in regression models fitted to the full data sets (figures S5B and S6B). High Shannon diversities occurred only when almost every bacterial species had a virulent phage predator (figures S5C and S6C). When systems under bottom-up and top-down control were analyzed separately, regression models reasonably described Shannon diversities (figures S5D,E and S6D,E). Systems with more phage species had higher Shannon diversities (table S4). The effect of temperate phages was larger when systems were under bottom-up control, but the effect of virulent phages was larger when systems were under top-down control.

**Fig. S1.**
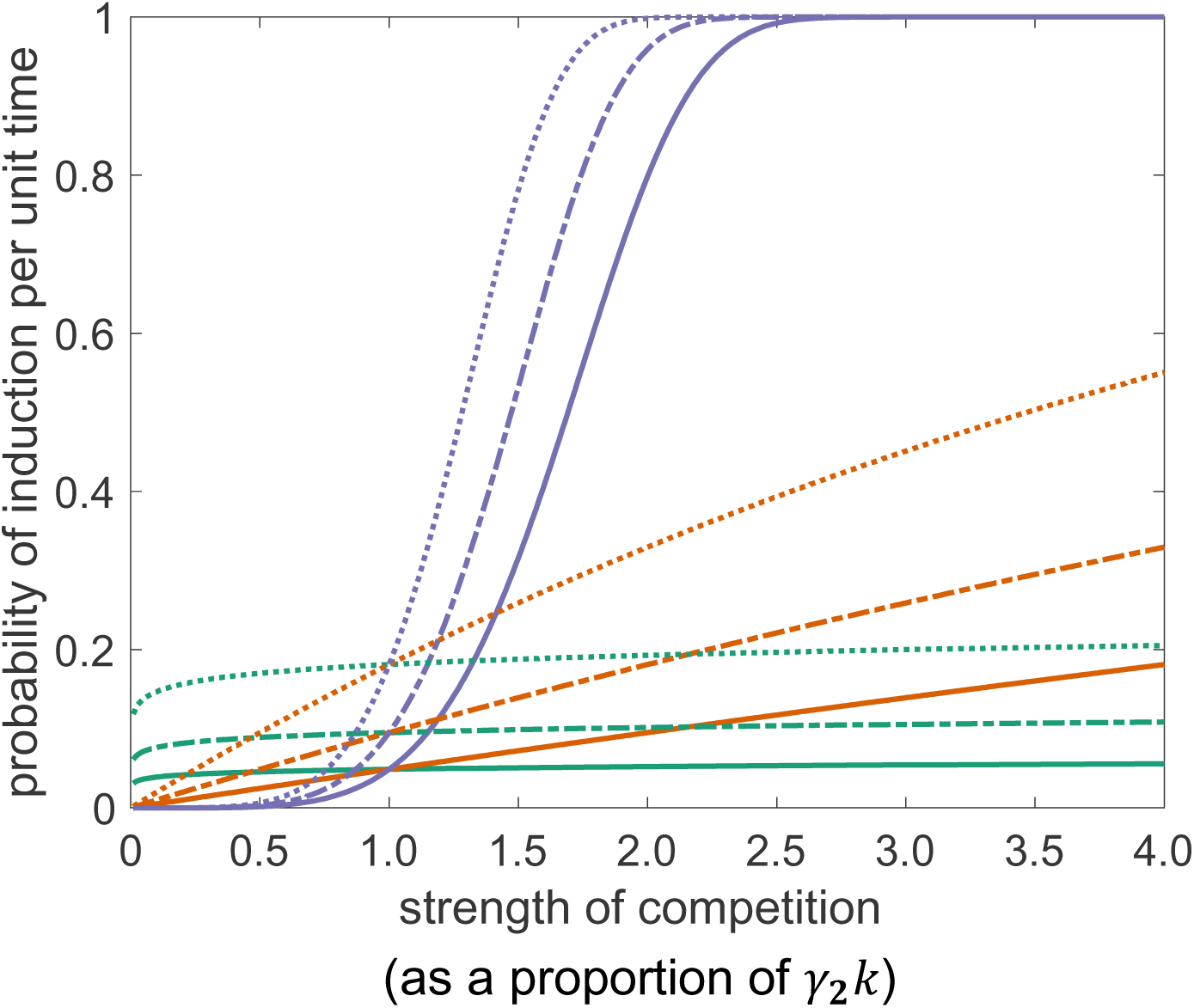
The parameterization competition-dependent induction in our model is capable of capturing a wide range of plausible induction functions. This figure shows the probability of induction per unit time as a function of the competition experienced by the lysogen. Solid lines show functions with 𝜸_𝟏_ = 𝟎. 𝟎𝟓, dashed lines show functions with 𝜸_𝟏_ = 𝟎. 𝟏, and dotted lines show functions with 𝜸_𝟏_ = 𝟎. 𝟐. **Green** lines show functions with 𝜸_𝟑_ = 𝟎. 𝟏, **brown** lines show functions with 𝜸_𝟑_ = 𝟏. 𝟎𝟏, and **purple** lines show functions with 𝜸_𝟑_ = 𝟓. 𝟎. The induction rate also depends on the parameter 𝜸_𝟐_. When 𝜸_𝟐_ is small, high induction rates are achieved at low proportions of carrying capacity, and when 𝜸_𝟐_ is large high induction rates are achieved only at high proportions of carrying capacity.

**Fig. S2.**
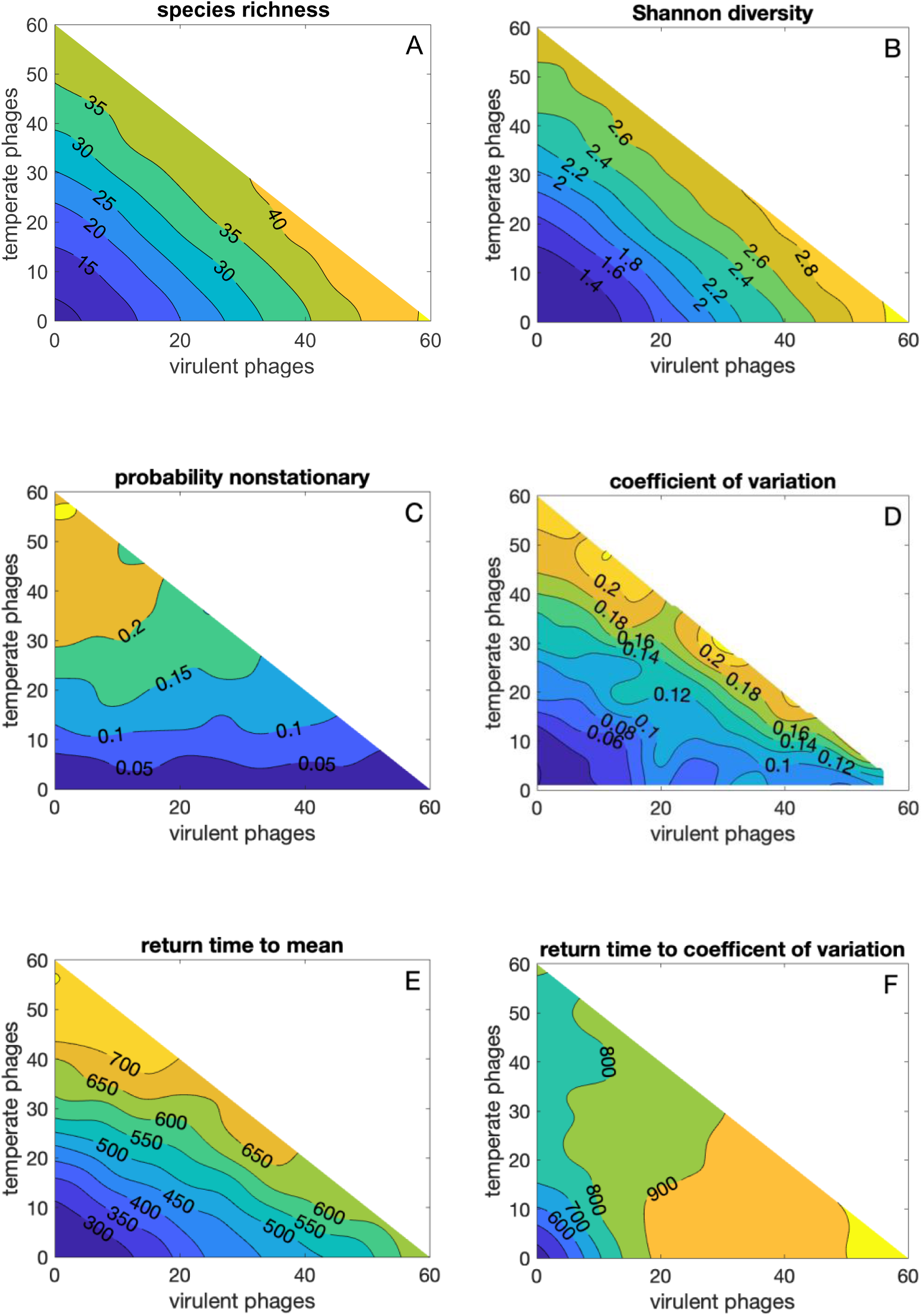
Diversity and stability measures as functions of the numbers of virulent and temperate generalist phage species in the species pool. Upper triangles are not used because systems were capped at 60 phage species. White areas in the lower triangle of (D) indicate that there were no nonstationary dynamics for those combinations of phage types.

**Fig. S3.**
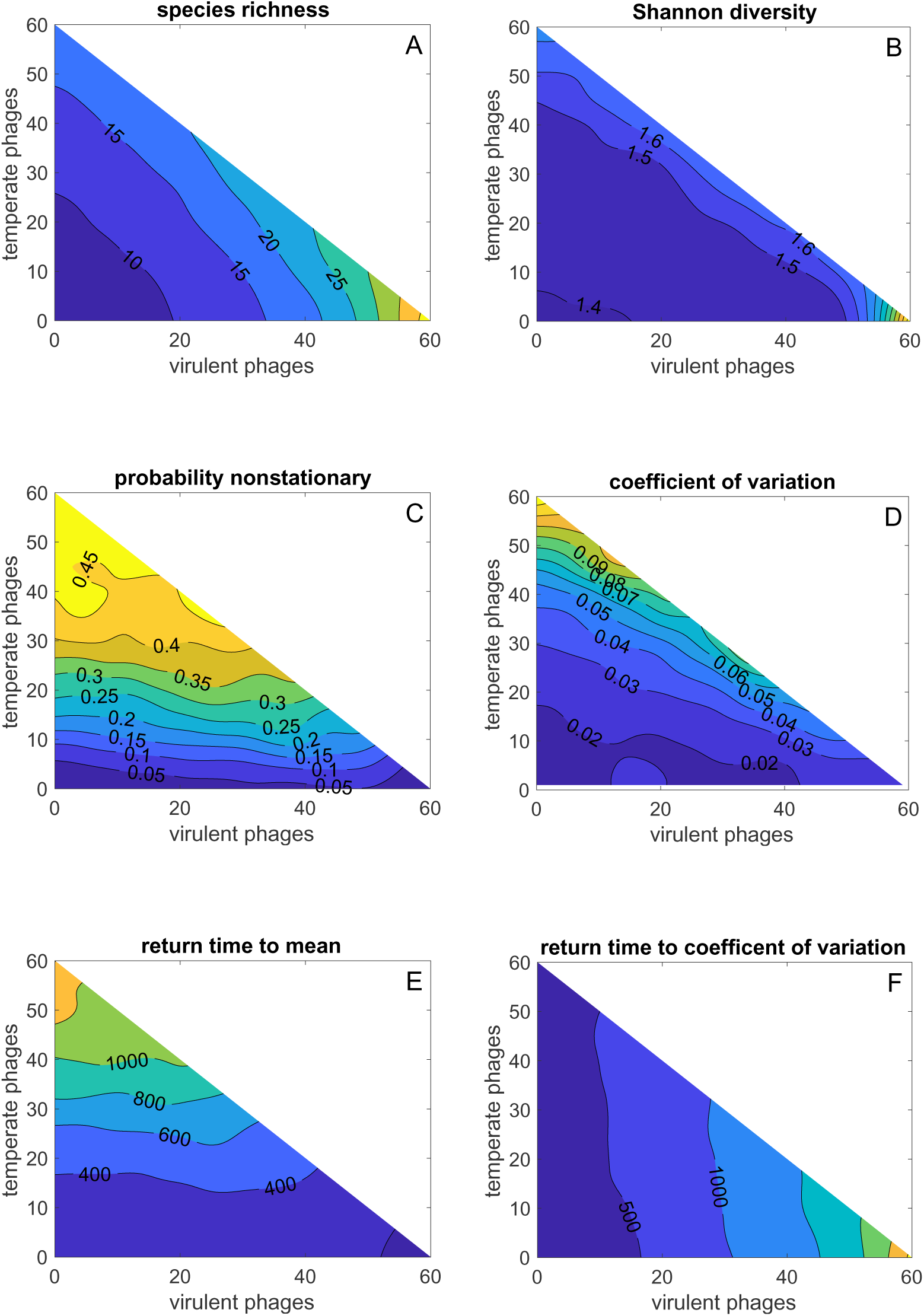
Diversity and stability measures as functions of the numbers of virulent and temperate phage species in the species pool when the probability of lysogeny and the induction rate were density dependent. Upper triangles are not used because systems were capped at 60 phage species. White areas of the lower triangle of (D) indicate that there were no nonstationary dynamics in the absence of temperate phages.

**Fig. S4.**
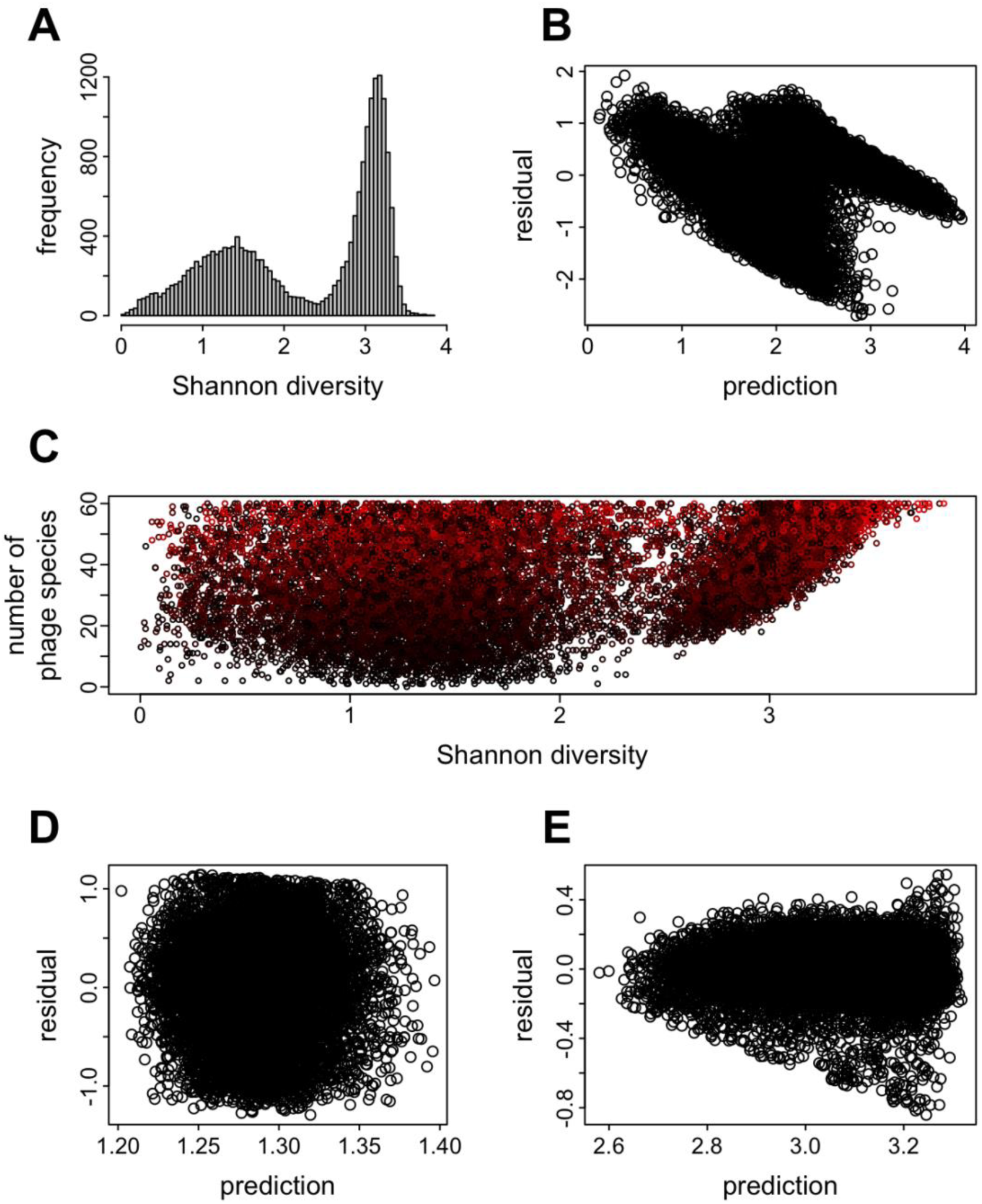
(A) Shannon diversities of bacterial communities in the mature state in 2 x 10^4^ simulations with generalist phages. (B) Residuals plotted against predicted Shannon diversities from regression models fitted to the same data. (C) Numbers of phage species in species pools for systems with different Shannon diversities of bacterial species in the mature state. Marker color indicates the number of virulent phage species (0:60 runs from black to red). (D,E) Residuals of regression models fitted to systems with bacterial Shannon diversities less than (D) or greater than (E) 2.4.

**Fig. S5.**
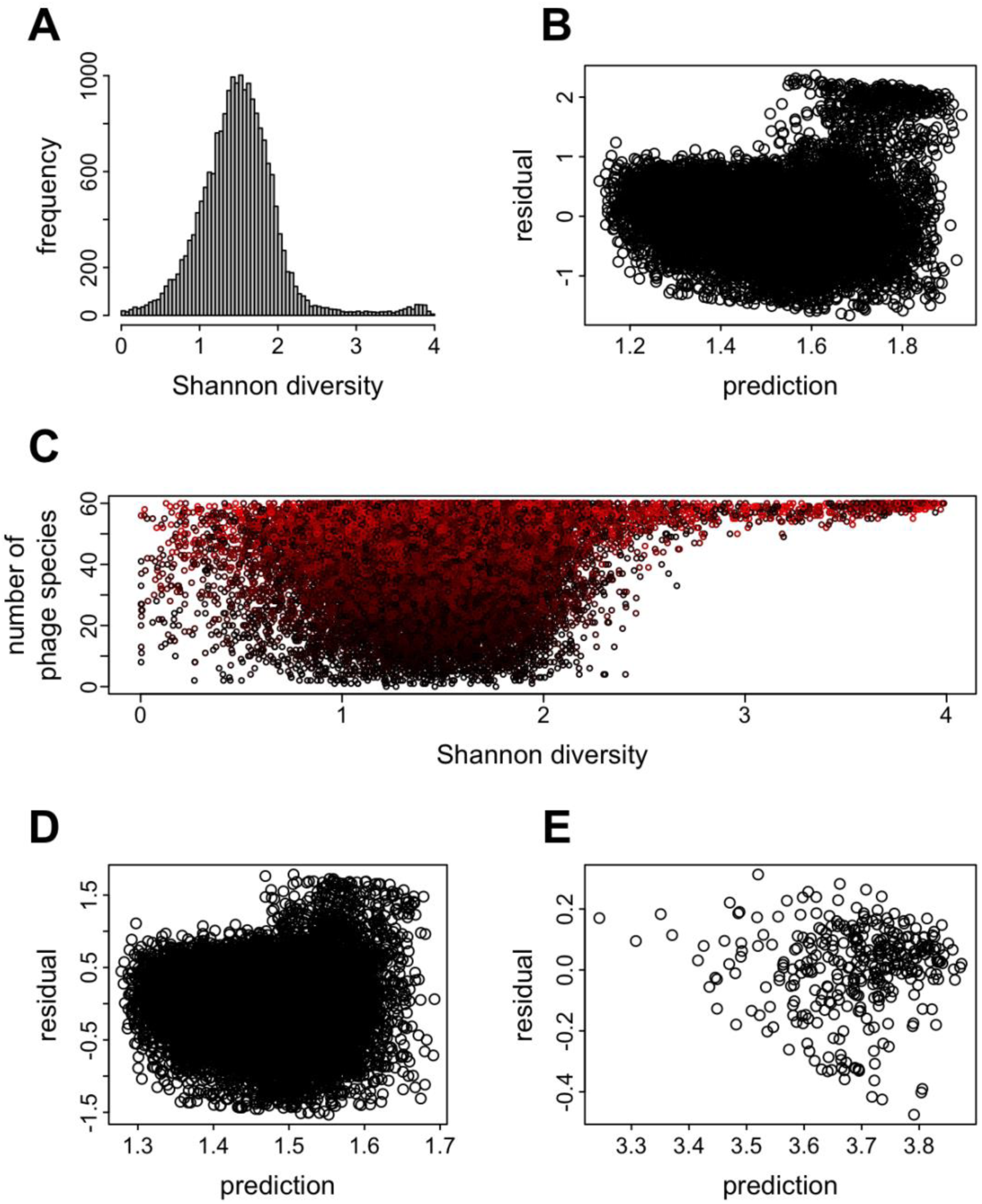
(A) Shannon diversities of bacterial communities in the mature state in 2 x 10^4^ simulations with specialist phages. (B) Residuals plotted against predicted Shannon diversities from regression models fitted to the same data. (C) Numbers of phage species in species pools for systems with different Shannon diversities of bacterial species in the mature state. Marker color indicates the number of virulent phage species (0:60 runs from black to red). (D,E) Residuals of regression models fitted to systems with bacterial Shannon diversities less than (D) or greater than (E) 3.3.

**Fig. S6.**
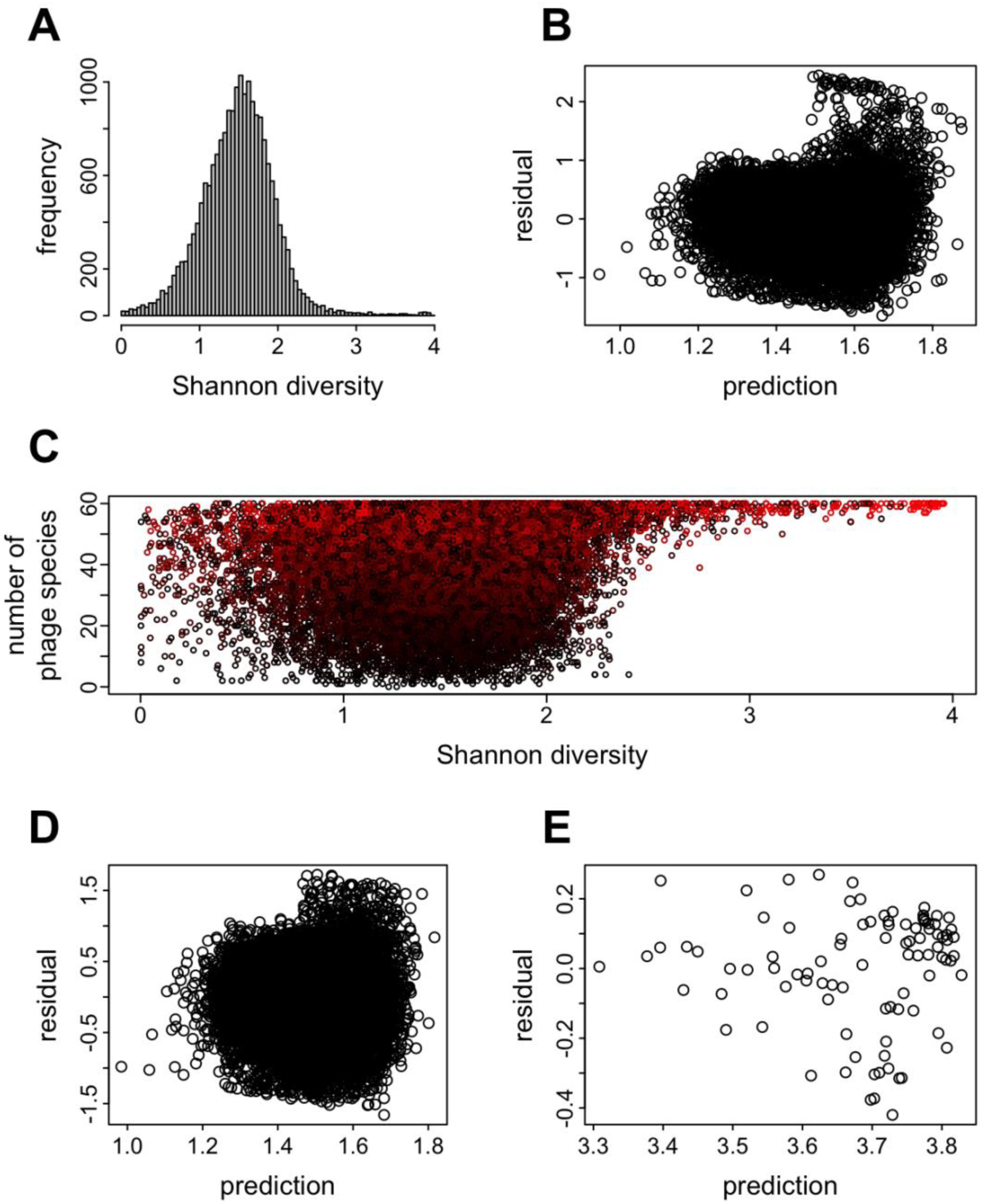
(A) Shannon diversities of bacterial communities in the mature state in 2 x 10^4^ simulations with specialist phages and density dependent probabilities of lysogeny and induction rates. (B) Residuals plotted against predicted Shannon diversities from regression models fitted to the same data. (C) Numbers of phage species in species pools for systems with different Shannon diversities of host species in the mature state. Marker color indicates the number of virulent phage species (0:60 runs from black to red). (D,E) Residuals of regression models fitted to systems with bacterial Shannon diversities less than (D) or greater than (E) 3.3.

**Table S1.**
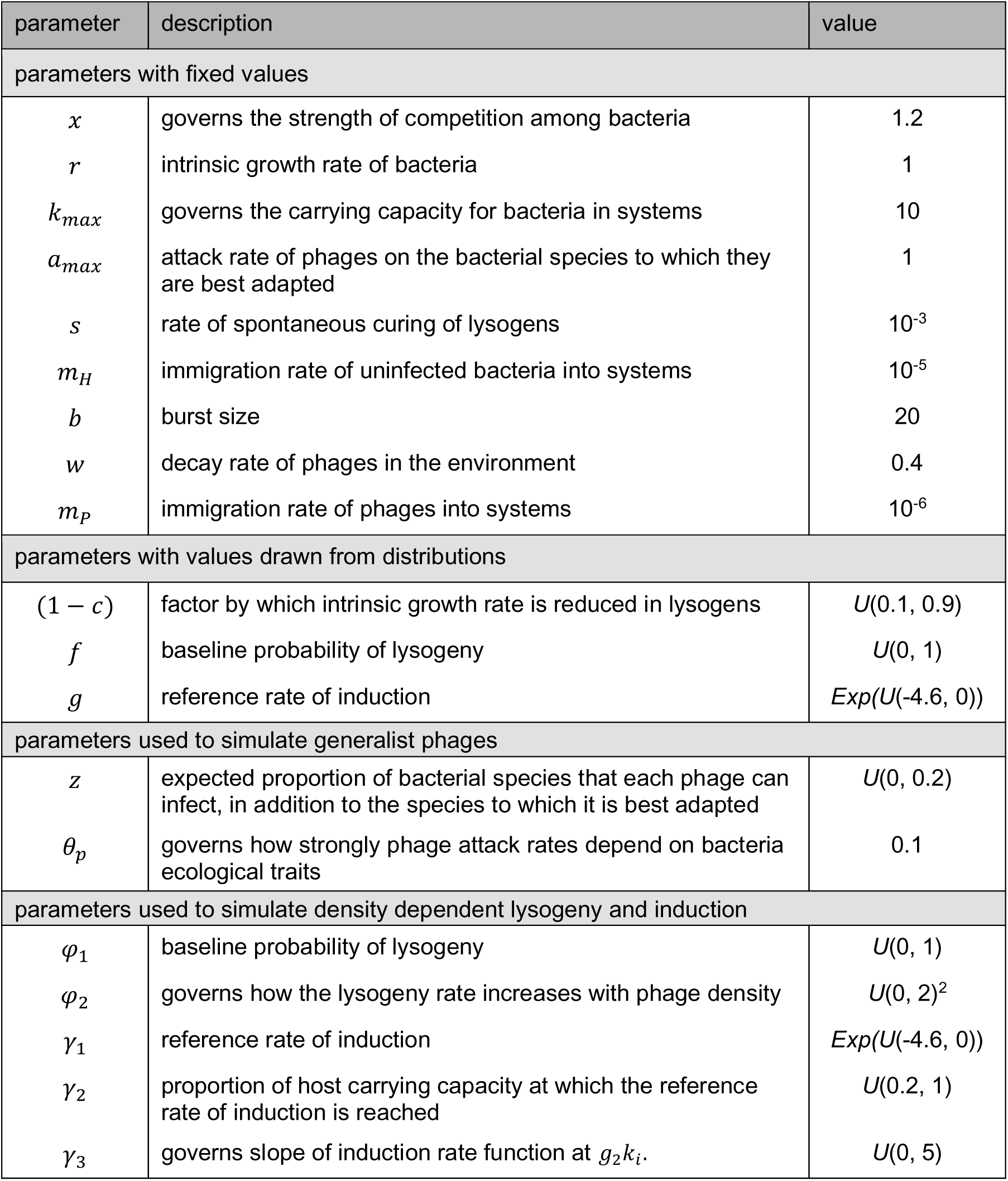
Parameters used in simulations.

**Table S2.** The effects of the numbers of virulent and temperate phage species and of the parameters that control lysogeny on the diversity and stability of bacterial communities when phages are generalists. Effects of phage species are reported as the change for each additional phage species in the species pool. The effects on the probability that a system is nonstationary are reported as the change in log odds ratios. The top (bottom) number in each cell indicates the effect of the predictor without (while) controlling for the diversity and variability of the bacterial community in the mature state. Nonsignificant effects are reported as ns. Parameters are set as in table S1.

| predictor | logged<br>species<br>richness | Shannon<br>diversity | probability<br>nonstationary | logged coefficient<br>of variation (CoV) | logged<br>return time<br>to mean | logged<br>return time<br>to CoV |
| --- | --- | --- | --- | --- | --- | --- |
| number of phage species |  |  |  |  |  |  |
| virulent | 0.0227 | 0.0357 | -0.0053<br>-0.0484 | 0.0428<br>ns | 0.0146<br>0.0043 | 0.0095<br>0.0021 |
| temperate | 0.0204 | 0.0307 | 0.0502<br>0.0149 | 0.0488<br>0.0090 | 0.0275<br>0.0129 | 0.0030<br>0.0013 |
| model parameters |  |  |  |  |  |  |
| cost of<br>lysogeny<br>(*c*) | 0.062 | 0.1038 | 2.4032<br>2.2694 | 1.0133<br>0.7786 | 0.3704<br>0.2174 | 0.1006<br>0.1807 |
| probability<br>of lysogeny<br>(*f*) | ns | ns | 1.8146<br>1.8167 | 0.5616<br>0.4903 | ns<br>ns | ns<br>ns |
| induction<br>rate<br>(*g*) | 0.1380 | 0.2744 | 3.7891<br>3.4933 | 1.4277<br>1.0377 | 0.6995<br>0.3297 | -0.1194<br>-0.0774 |
| proportion<br>of bacterial<br>species<br>each<br>phage can<br>infect (*z*) | 4.1632 | 8.117 | -12.3716<br>-21.5414 | 11.0184<br>1.9256 | 0.6383<br>-0.9245 | -1.4800<br>-3.3790 |
| (1 – *f*) x<br>number of<br>temperate<br>phage<br>species | ns | ns | ns<br>ns | ns<br>ns | -0.0139<br>-0.0090 | ns<br>-0.0039 |
| emergent properties of systems |  |  |  |  |  |  |
| species<br>richness |  |  | 0.0824 | 0.0321 | 0.0179 | 0.0070 |
| Shannon<br>diversity |  |  | -0.2873 | 0.6082 | -0.0709 | 0.0911 |
| coefficient<br>of variation<br>(CoV) |  |  |  |  | 3.5890 | -3.0070 |

**Table S3.** The effects of the numbers of virulent and temperate phage species and of the parameters that control lysogeny on the diversity and stability of bacterial communities when phages are specialists and lysogeny and induction are density dependent. Effects of phage species are reported as the change for each additional phage species in the species pool. The effects on the probability that a system is nonstationary are reported as the change in log odds ratios. The top (bottom) number in each cell indicates the effect of the predictor without (while) controlling for the diversity and variability of the bacterial community in the mature state. Nonsignificant effects are reported as ns. Parameters are set as in table S1.

| predictor | logged species richness | Shannon diversity | probability nonstationary | logged coefficient of variation (CoV) | logged return time to mean | logged return time to CoV |
| --- | --- | --- | --- | --- | --- | --- |
| number of phage species |  |  |  |  |  |  |
| virulent | 0.0306 | 0.0053 | 0.0139<br>-0.0923 | 0.0168<br>-0.0182 | -0.0036<br>-0.0232 | 0.0458<br>0.0334 |
| temperate | 0.0196 | 0.0083 | 0.1009<br>0.0645 | 0.0548<br>0.0270 | 0.0402<br>0.0236 | 0.0055<br>0.0045 |
| model parameters |  |  |  |  |  |  |
| $c$ | 0.1980 | 0.1370 | 0.9109<br>0.4106 | 0.5008<br>0.2807 | 0.2803<br>0.0877 | -0.0426<br>-0.0547 |
| $\varphi_1$ | 0.0226 | -0.1169 | 1.6101<br>1.4295 | -0.3615<br>-0.2085 | 0.2383<br>0.1324 | ns<br>-0.0637 |
| $\varphi_2$ | ns | ns | 0.1577<br>0.1873 | 0.0518<br>0.0568 | 0.0389<br>0.0290 | -0.0095<br>ns |
| $\gamma_1$ | 0.4462 | 0.1540 | 5.1099<br>4.3206 | 0.8095<br>0.4183 | 1.0530<br>0.3955 | ns<br>0.1002 |
| $\gamma_2$ | -0.3916 | -0.1239 | -4.4290<br>-3.5380 | -1.0300<br>-0.4744 | -0.6660<br>-0.1809 | -0.0807<br>-0.1034 |
| $\gamma_3$ | 0.0090 | 0.0062 | 0.2058<br>0.1983 | -0.0620<br>-0.0458 | ns<br>ns | 0.0322<br>0.0237 |
| observed lysis rate x number of temperate phage species | 0.0032 | -0.0185 | -0.0985<br>-0.1731 | -0.0983<br>-0.0950 | -0.0411<br>-0.0389 | 0.0269<br>0.0182 |
| emergent properties of systems |  |  |  |  |  |  |
| species richness |  |  | 0.2107 | 0.0624 | 0.0434 | 0.0255 |
| Shannon diversity |  |  | -1.2641 | -0.0722 | -0.5288 | 0.1825 |
| coefficient of variation (CoV) |  |  |  |  | 7.6740 | -6.2930 |

**Table S4.** The effects of the numbers of generalist virulent and temperate phage species on Shannon diversities in bacterial communities under bottom-up and top-down control. Effect sizes are per phage species in the species pool.

| simulation set | control regime | Shannon diversities | effect of virulent phage species | effect of temperate phage species |
| --- | --- | --- | --- | --- |
| generalist phages | bottom-up | <2.4 | 0.0009 | 0.0032 |
|  | top-down | >2.4 | 0.0140 | 0.0118 |
| specialist phages | bottom-up | <3.3 | 0.0036 | 0.0055 |
|  | top-down | >3.3 | 0.0515 | 0.0488 |
| specialist phages with density dependent lysogeny and induction | bottom-up | <3.3 | 0.0038 | 0.0079 |
|  | top-down | >3.3 | 0.0298 | 0.0288 |

