## Supplementary material for "Lysogeny destabilizes computationally simulated microbiomes": a.Codes and data for Gilman et al 2023.zip: analyse_results_dd.R

```
library(lme4)
library(lmerTest)

#load data set
data=read.csv("spec_dd_full.csv")

#assign column names
colnames(data)=c("num.phage.species","prop.host.strains.vulnerable","nonspecialist.interactions",
                 "phi1","prob.phage.temperate","num.temperate.phages",
                 "gamma2","gamma3","victim.richness","victim.shannon",
                 "victim.COV","rt.mean","rt.range","rt.stds",
                 "sim.seed","lyap","phi2","mature.lysis.rate",
                 "gamma1","nu","c","rt.Aitch","rt.CoV")

data$lysogeny.cost=1-data$c
data$num.virulent.phages=data$num.phage.species-data$num.temperate.phages
data$lysis.rate=1-data$mature.lysis.rate
data$lysis.rate[which(is.na(data$lysis.rate))]=0

#study victim species richness (results in table S3)
data3=data[c(1,3*(1:(nrow(data)/3))),]
vr.glm=glm(victim.richness~num.virulent.phages+num.temperate.phages+lysogeny.cost+phi1+phi2+gamma1+gamma2+gamma3+lysis.rate:num.temperate.phages,data=data3,family="poisson")
summary(vr.glm)
hist(resid(vr.glm),100)
plot(predict(vr.glm),resid(vr.glm))

#study victim Shannon diversity (results in table S3)
vs.lm=lm(victim.shannon~num.virulent.phages+num.temperate.phages+lysogeny.cost+phi1+phi2+gamma1+gamma2+gamma3+lysis.rate:num.temperate.phages,data=data3)
summary(vs.lm)
hist(resid(vs.lm))
plot(predict(vs.lm),resid(vs.lm),xlab="prediction",ylab="residual",main="")

#split between modes
data3.p1=data3[which(data3$victim.shannon<3.3),]
data3.p2=data3[which(data3$victim.shannon>3.3),]

#study first mode (results in table S4)
vsp1.lm=lm(victim.shannon~num.virulent.phages+num.temperate.phages+lysogeny.cost+phi1+gamma1+lysis.rate:num.temperate.phages,data=data3.p1)
summary(vsp1.lm)
hist(resid(vsp1.lm))

#study second mode (results in table S4)
vsp2.lm=lm(victim.shannon~num.virulent.phages+num.temperate.phages+lysogeny.cost+phi1+gamma1+lysis.rate:num.temperate.phages,data=data3.p2)
summary(vsp2.lm)
hist(resid(vsp2.lm))

#study probability that the victim populations are intrinsically nonstationary (results in table S3)
non.stat=(data3$victim.COV>0.01)
vns.glm=glm(non.stat~num.virulent.phages+num.temperate.phages+lysogeny.cost+phi1+phi2+gamma1+gamma2+gamma3+lysis.rate:num.temperate.phages,data=data3,family="binomial")
summary(vns.glm)

#study probability that the victim populations are intrinsically nonstationary controlling for diversity (results in table S3)
vns.glm=glm(non.stat~num.virulent.phages+num.temperate.phages+lysogeny.cost+phi1+phi2+gamma1+gamma2+gamma3+lysis.rate:num.temperate.phages+victim.richness+victim.shannon,data=data3,family="binomial")
summary(vns.glm)

#study victim CoV when non-stationary (results in table S3)
data3.cov=data3[which(data3$victim.COV>0.01),]
vcov.lm=lm(log(victim.COV)~num.virulent.phages+num.temperate.phages+lysogeny.cost+phi1+phi2+gamma1+gamma2+gamma3+lysis.rate:num.temperate.phages,data=data3.cov)
summary(vcov.lm)
hist(resid(vcov.lm))

#study victim CoV when non-stationary controlling for diversity (results in table S3)
vcov.lm=lm(log(victim.COV)~num.virulent.phages+num.temperate.phages+lysogeny.cost+phi1+phi2+gamma1+gamma2+gamma3+lysis.rate:num.temperate.phages+victim.richness+victim.shannon,data=data3.cov)
summary(vcov.lm)
hist(resid(vcov.lm))

#study return time to mean density (results in table S3)
data$group=as.factor(data$phi1)
data$rt.mean[which(data$rt.mean>5000)]=5000
vrtm.lm=lmer(log(rt.mean)~num.virulent.phages+num.temperate.phages+lysogeny.cost+phi1+phi2+gamma1+gamma2+gamma3+lysis.rate:num.temperate.phages+(1|group),data=data)
summary(vrtm.lm)
hist(resid(vrtm.lm),100)
plot(predict(vrtm.lm),resid(vrtm.lm))

#study return time to mean densigty controlling for diversity and dynamics (results in table S3)
vrtm.lm=lmer(log(rt.mean)~num.virulent.phages+num.temperate.phages+lysogeny.cost+phi1+phi2+gamma1+gamma2+gamma3+lysis.rate:num.temperate.phages+victim.richness+victim.shannon+victim.COV+(1|group),data=data)
summary(vrtm.lm)
hist(resid(vrtm.lm),100)
plot(predict(vrtm.lm),resid(vrtm.lm))

#study return time to CoV (results in table S3)
data$rt.CoV[which(data$rt.CoV>5000)]=5000
vrtm.lm=lmer(log(rt.CoV)~num.virulent.phages+num.temperate.phages+lysogeny.cost+phi1+phi2+gamma1+gamma2+gamma3+lysis.rate:num.temperate.phages+(1|group),data=data)
summary(vrtm.lm)
hist(resid(vrtm.lm),100)
plot(predict(vrtm.lm),resid(vrtm.lm))

#study return time to CoV in just stationary systems
data.stat=data[which(data$victim.COV>10^-2),]
vrtm.lm=lmer(log(rt.CoV)~num.virulent.phages+num.temperate.phages+lysogeny.cost+phi1+gamma1+lysis.rate:num.temperate.phages+(1|group),data=data.stat)
summary(vrtm.lm)
hist(resid(vrtm.lm),100)
plot(predict(vrtm.lm),resid(vrtm.lm))

#study return time to CoV controlling for diversity and dynamics (results in table S3)
vrtm.lm=lmer(log(rt.CoV)~num.virulent.phages+num.temperate.phages+lysogeny.cost+phi1+phi2+gamma1+gamma2+gamma3+lysis.rate:num.temperate.phages+victim.richness+victim.shannon+victim.COV+(1|group),data=data)
summary(vrtm.lm)
hist(resid(vrtm.lm),100)
plot(predict(vrtm.lm),resid(vrtm.lm))


#make figure S6
par(cex=.85, mai=c(0.1,0.1,0.2,0.1))

#histogram
par(fig=c(0.1,0.45,0.7,0.95))
hist(data3$victim.shannon,100,main="",axes=F)
mtext("Shannon diversity",1,line=1.75)
mtext("frequency",2,line=1.75)
axis(1,tck=-.02,mgp=c(3,.25,0))
axis(2,tck=-.02,mgp=c(3,.25,0))

#residuals
par(fig=c(0.55,0.95,0.7,0.95), new=TRUE)
plot(predict(vs.lm),resid(vs.lm),main="",axes=F,frame=T)
mtext("prediction",1,line=1.75)
mtext("residual",2,line=1.75)
axis(1,tck=-.02,mgp=c(3,.25,0))
axis(2,tck=-.02,mgp=c(3,.25,0))

#scatter
par(fig=c(0.1,0.95,0.4,0.6), new=TRUE)
plot(data3$victim.shannon,data3$num.phage.species,col=rgb(data3$num.virulent.phages/60,0,0),cex=.5,xlab="Shannon diversity",ylab="number of phage species",mgp=c(3,.25,0),axes=F,frame=T)
axis(1,tck=-.04,mgp=c(3,.25,0))
axis(2,tck=-.04,mgp=c(3,.25,0))
mtext("Shannon diversity",1,line=1.75)
mtext("number of\nphage species",2,line=1.75)

#residuals (low)
par(fig=c(0.1,0.45,0.05,0.3), new=TRUE)
plot(predict(vsp1.lm),resid(vsp1.lm),main="",axes=F,frame=T)
mtext("prediction",1,line=1.75)
mtext("residual",2,line=1.75)
axis(1,tck=-.02,mgp=c(3,.25,0))
axis(2,tck=-.02,mgp=c(3,.25,0))

#residuals (high)
par(fig=c(0.55,0.95,0.05,0.3), new=TRUE)
plot(predict(vsp2.lm),resid(vsp2.lm),main="",axes=F,frame=T)
mtext("prediction",1,line=1.75)
mtext("residual",2,line=1.75)
axis(1,tck=-.02,mgp=c(3,.25,0))
axis(2,tck=-.02,mgp=c(3,.25,0))

# title(outer=T,adj=.04,main="A",cex.main=2,line=-2.5)
# title(outer=T,adj=.54,main="B",cex.main=2,line=-2.5)
# title(outer=T,adj=.04,main="C",cex.main=2,line=-18)
# title(outer=T,adj=.04,main="D",cex.main=2,line=-32)
# title(outer=T,adj=.54,main="E",cex.main=2,line=-32)
```
