## Supplementary material for "Lysogeny destabilizes computationally simulated microbiomes": a.Codes and data for Gilman et al 2023.zip: get_dVP_nm.m

```
function [dVP]=get_dVP_nm(X,Alpha,K,r,mh,mp,A,b,d,w,f1,f2,g1,x,g2,g3,c,s,lys_apt)

Pv=size(A,1);

%create objects for equations
H=X(1:Pv);
P=X(1+Pv:2*Pv);
L=X(1+2*Pv:3*Pv);
%Lambda=(1-(1-f1)*exp(-f2*P)).*lys_apt;
Lambda=(f1+(1-f1)*(P./(P+f2^-1))).*lys_apt;
Omega=A.*(1-eye(Pv))+diag(diag(A).*(1-Lambda));
Upsilon=A.*(1-eye(Pv));

dV=r*(1-Alpha*(H+L)./K).*H...  %population growth 
    -A*P.*H... %loss due to virus attack 
    +s*L...    %spontenious curing of lysogens 
    +mh... %immigration 
    -w*H; %washout (not used, w=0);

dP=(b-1)*(Omega'*H).*P... %viruses created by direct lysis of non-lysogens
    +(b-1)*(Upsilon'*L).*P... %new viruses from attack on susceptible lysogens
    -((A-Omega)'*H).*P... %virus lost in creation of lysogens
    +b*g1*((Alpha*(H+L)./(g2*K)).^g3).*L...  %new virus by induction 
    -d*P... %death or viral wash out 
    +mp; %immigration of viruses
    
dL=c*r*(1-Alpha*(H+L)./K).*L...     %population growth
    +((A-Omega)'*H).*P...    %creation of lysogens
    -g1*((Alpha*(H+L)./(g2*K)).^g3).*L... %density dependent induction
    -Upsilon*P.*L...  %lysogens destroyed by other viruses
    -s*L... %spontaneous "curing" of lysogens
    -w*L; %washout (not used, w=0);
    
dVP=real([dV;dP;dL]);
```
