## Supplementary material for "Lysogeny destabilizes computationally simulated microbiomes": a.Codes and data for Gilman et al 2023.zip: hello_chaos_nm.m

```
function [lyap]=hello_chaos_nm(X,Pv,dVP,i)

options_fine=odeset('NonNegative',[1:(3*Pv)],'RelTol',1e-6,'AbsTol',1e-8);
[t,sol] = ode15s(dVP,[0:0.5:1000],X,options_fine);

%outname=strcat('lyap_dat_',string(i),'.csv');
%writematrix(sol,outname)

%lyap=42;
lyap=lyapunovExponent(sol);
```
