## Supplementary material for "Lysogeny destabilizes computationally simulated microbiomes": a.Codes and data for Gilman et al 2023.zip: make_surfs.m

```
%read in the appropriate data
od=readmatrix('spec_no_dd_full.csv');

%OPTIONAL: Ignore systems that are or are not stationary. If this is
%commented, then the plot of CoV will look at the probability that the
%mature state is non-stationary. If it is uncommented, the plot will look
%at the magnitude of the CoV
%od(od(:,11)<0.01,:)=[];

%choose results to plot (9 species richness, 10 Shannon index, 11
%Coefficient of Variation, 12 return time to mean density, 23 return to CoV)
ri=23;

%shrink od for other than return times
if ((ri<12 | ri>14) & (ri<22 | ri>23))
    od=od(3:3:size(od,1),:);
else
    %if return times are infinite, set them to maximum
    od(od(:,ri)==Inf,ri)=5000;
    %average the logged return times to make sample size manageable
    od(:,ri)=log(od(:,ri));
    sten=sparse(size(od,1)/3,size(od,1));
    stenr=size(sten,1);
    steni=1:stenr;
    sten((steni-1)*3*stenr+steni)=1;
    sten((steni-1)*3*stenr+stenr+steni)=1;
    sten((steni-1)*3*stenr+2*stenr+steni)=1;
    rvect=(sten*od(:,ri))/3;
    od=od(3:3:size(od,1),:);
    od(:,ri)=rvect;
end

%create axes
p1=od(:,1)-od(:,6); %non-lysogenic viruses
p2=od(:,6);
r=od(:,ri);

% %remove infinites and set -Inf to minimum observed value
% minr=min(r(r>-Inf));
% r(r==-Inf)=minr;
% p1(r==Inf)=[];
% p2(r==Inf)=[];
% r(r==Inf)=[];

%transform speices counts
if (ri==9)
    r=log(r);
%make stationarity categorical    
elseif (ri==11 & min(r)<0.001)
    r=(r>0.01);
end


%set range
res=100;

%get p1 and p2 scaled to range
p1s=(p1-min(p1))/(max(p1)-min(p1));
p2s=(p2-min(p2))/(max(p2)-min(p2));

%compute weighted means with std sig
sig=0.05;
%sig=0.1;
gdim1=((0:res)'/res)*ones(1,res+1);
gdim2=ones(res+1,1)*((0:res)/res);
%get the distance of each scaled observation from the each grid point,
%where the grid points are in rows and the observations are in columns
p1dist=(p1s*ones(1,(res+1)^2)-ones(length(p1s),1)*gdim1(1:(res+1)^2))';
p2dist=(p2s*ones(1,(res+1)^2)-ones(length(p2s),1)*gdim2(1:(res+1)^2))';
%now, for every grid point, compute the weight of each observation
obs_weight=exp(-(p1dist.^2+p2dist.^2)/(2*sig^2));
%finally, compute the weighted mean observed value for each grid point
weighted_total_obs=obs_weight*r;
total_weight=obs_weight*ones(size(r));
r_on_grid=weighted_total_obs./total_weight;

%set min weight for plotting
r_on_grid(find(total_weight<1))=0/0;

%undo logging where necessary
if (ri==9 | ri==12 | ri==23)
    r_on_grid=exp(r_on_grid);
end

%create X, Y, Z matrices
x=min(p1):(max(p1)-min(p1))/res:max(p1);
y=min(p2):(max(p2)-min(p2))/res:max(p2);
X=x'*ones(1,res+1);
Y=ones(res+1,1)*y;
Z=X;
Z(1:(res+1)^2)=r_on_grid;

Z(X+Y>60)=0/0;

f=figure(1)
if ri==16
    [c,h]=contourf(X,Y,Z,.0:.1:.6,'ShowText','on');
    axis([1 60 1 60])
elseif ri==13
    [c,h]=contourf(X,Y,Z,200:200:2400,'ShowText','on');
else
    [c,h]=contourf(X,Y,Z,'ShowText','on');
end
xlim([0 60])
ylim([0 60])
clabel(c,h,'FontSize',15)
set(gca,'FontSize',15)
xlabel('virulent phages')
ylabel('temperate phages')
if ri==16
    title('Lyapunov exponent')
elseif ri==9
    title('species richness')
elseif ri==10
    title('Shannon diversity')
elseif ri==11
    if min(r)<0.001
        title('probability nonstationary')
    else
        title('coefficient of variation')
    end
elseif ri==12
    title('return time to mean')
elseif ri==13
    title('return time to range')
elseif ri==22
    title('return time to Aitchison distance')
elseif ri==23
    title('return time to coefficent of variation')
end

text(56,56,"B","FontSize",20)
f.Position=[360 402 371 296]

mean_response=exp(mean(r))
```
