## Supplementary material for "Lysogeny destabilizes computationally simulated microbiomes": a.Codes and data for Gilman et al 2023.zip: microbiome_simulator.m

```
%%%%%%%%%%
%This code runs the simulations described in Gilman et al. 2023
%Code by RTG, Levenshulme, last edit: 28 August 2023
%%%%%%%%%%

%Set seed number to make simulations replicable. In the paper we report
%results for seed numbers 0 to 19,999.
seednum = 0;

%Set parameters used in the paper.
%parameters that describe bacteria
Pv=60;      %number of bacterial species in the species pool 
nv=4;       %sets numnber of dimensions in ecological trait space for competition among bacteria
cv=1.2;     %scales competition among bacteria (corresponds to x in table S1)
Kmax=10;    %scales carrying capacity for bacteria
r=1;        %intrinsic bacterial population growth rate
mh=10^-5;   %immigration rate of bacteria into system
w=0;        %washout rate (not used)
%parameters that describe phages
np=2;       %number of ecological dimensions determing predation rate
att=1;      %maximum attack rate of phages on bacteria
cp=10;      %governs ecological breadth of bacterial species phages can attack (corresponds to 1/theta in the paper)
b=20;       %the burst size
d=.4;       %death rate for phages (corresponds to w in the paper)
x=0;        %rate at which phages are destroyed in attacks on lysogens (not used)
s=0.001;    %rate at which lysogens spontaneously "heal" to become uninfected hosts
%parameters controlling perturbations from mature states of systems
xi=3;       %number of perturbations to study
kick_size=0.5;  %magnitude of perturbations

%If our system has only 1 phage, we only need to study it when it has 0 or 1
%temperate phages. If it has 60 phages, then we need to study
%it when it has 0, 1, ..., 59, or 60 temperate phages. So, we need to
%study more systems with 60 phages than with 1. Pp_tool is a vector where
%entry i is the relative frequency of systems with fewer than i phage
%species. We will use when randomly creating systems later.
Pp_tool=cumsum((1:Pv+1)/sum(1:Pv+1));

%Create a matrix to hold the output data
out_dat=[];

for i=(seednum:(seednum+9))  %how many similations to run
    
    ci=i;                   %to track the run
    rng(ci);                %to make the runs repeatable
    
    %parameter values that vary among simulations
    Pp=sum(rand>Pp_tool);   %number of phage species in simulation
    z=0;                    %proportion of host strains that each phages can infect (in addition to the one on which is sopecialises)
    f1=rand(1);             %baseline lysogeny rate (phi1 in the paper)
    f2=0;                   %controls density dependence of the lysogeny rate (phi2 in the paper)
    lp=rand(1);             %proportion of phage species in the simulation that are temperate
    g2=0.2+0.8*rand(1);     %proportion of host carrying capacity at which the baseline induction rate is reached (gamma 2 in the paper)
    g3=0;                   %governs slope of the induction rate function (gamma3 in the paper)
    g1=exp(-4.6*rand(1));   %baseline induction rate (gamma1 in paper)
    c=0.1+0.8*rand(1);      %governs cost of lysogeny to reproduction (c in paper)
    
    %Create the system of ODEs ans simulate it until it reaches its mature
    %state
    [Xfinal,Xeq,XVfinal,XVeq,XVmin,XVmax,XVstd,XVclr,tot_victim_sr,tot_victim_shan,tot_victim_wcov,A,K,pred_apt,mp,tu,lys_apt,dVP,mean_lys]=v_and_p_func_nm(Pv,Pp,nv,cv,Kmax,r,mh,w,np,att,z,cp,b,d,f1,f2,g1,x,g2,g3,c,s,lp);
    
    %Estimates Lyapunov exponent of system in mature state
    if tot_victim_wcov>(10^-2)
        lyap=hello_chaos_nm(Xfinal(end,:),Pv,dVP,i);
    else
        lyap=-Inf;
    end
    
    %Studies return times to mature states
    [rt_vect,tn,soln]=qcc_func_nm(i,xi,kick_size,Xfinal(end,:),Pv,tu,XVeq,XVmax,XVmin,XVstd,XVclr,tot_victim_wcov,dVP);

    %Stores results of simulation
    %[Pp,z,pred-prey interaction count,f1,lp,lysogenic virus count, [1-6]...
    %g2,g3,richness of victims,Shannon diversity of victims,weighted CoV of victims, [7-11]...
    %return time for means, return time for bounds,return time for stds, [12-14]...
    %simulation seed, lyapunov exponent,f2,mean lysis rate,g1,nu,c [15-21]]
    
    out_dat=[out_dat;[ones(xi,1)*[Pp,z,sum(sum(sign(pred_apt-eye(Pv))>0)),f1,lp,sum(lys_apt),...
        g2,g3,tot_victim_sr,tot_victim_shan,tot_victim_wcov],rt_vect(:,1:3),ones(xi,1)*[ci,lyap,f2,mean_lys,g1,0,c],rt_vect(:,4:5)]];
    
    %Saves results to a csv 
    if rem(i,10)==0
        %Uncommment to save results every 10 runs in case of 
%         outname=strcat('mic_sim_spec_no_dd_',string(seednum),'.csv');
%         writematrix(out_dat,outname)
    end
    
end

%Saves results to a csv 
% outname=strcat('mic_sim_spec_no_dd_',string(seednum),'.csv');
% writematrix(out_dat,outname)
```
