## Supplementary material for "Lysogeny destabilizes computationally simulated microbiomes": a.Codes and data for Gilman et al 2023.zip: qcc_func_nm.m

```
function [rt_vect,tn,soln]=qcc_func_nm(ci,xi,kick_size,X2kick,Pv,tu,XVeq,XVmax,XVmin,XVstd,XVclr,tot_victim_wcov,dVP)

rt_vect=[];
fstates=[];

options_fine=odeset('NonNegative',[1:(3*Pv)],'RelTol',1e-6,'AbsTol',1e-8);

for ii=1:xi

    %kick Xeq delivered from v_and_p_func_nm.m
    X_k=X2kick.*exp(kick_size*sign(randn(size(X2kick)))); 

    %solve from the kicked state
    X=X_k';
    [tn,soln] = ode15s(dVP,[0:5000],X,options_fine);
% %     Xfinal_n=soln(tn>=tn(end)-tu,:);    %needed?
% %     Xeq_n=mean(Xfinal_n,1);             %needed?

    %check the last time any victim was away from its pre-kick state by more
    %than th1 OR cycles were more than th2 larger than in the final state
    %from the original simulation
    th1=0.01;
    th2=0.05;
    tu_n=200;   %for how long do we examine in the post-kick state?

    ret_time=[0,0,0,0,0];
    i=sum(tn<2); %so there are at least 2 time steps
    while min(ret_time)==0 %find return times for means, CoVs, and ranges

        XVfinal_n=soln(((tn<=tn(i))&(tn>tn(i)-tu_n)),1:Pv)...
            +soln(((tn<=tn(i))&(tn>tn(i)-tu_n)),1+2*Pv:3*Pv);
        
        %compute mean over time
        XVeq_n=mean(XVfinal_n,1);  
        tnow=tn(((tn<=tn(i))&(tn>tn(i)-tu_n)));

        XVmin_n=min(XVfinal_n,[],1);
        XVmax_n=max(XVfinal_n,[],1);
        XVstd_n=std(XVfinal_n,[],1);

        XVclr_n=log(XVeq_n)-mean(log(XVeq_n));
        XVwcov_n=sum(XVstd_n)/sum(mean(XVfinal_n));
   
        if (max(abs(log(XVeq_n./XVeq)).*(XVeq_n>10^-4*sum(XVeq_n)))<th1...
                & ret_time(1)==0)
            ret_time(1)=(min(tnow)+max(tnow))/2;
            %population means must be within th1 of the pre-kick
            %value and greater than 10^-4 proportionally
        end
        if (sqrt(sum((XVclr-XVclr_n).^2))<0.01 & ret_time(4)==0)
            ret_time(4)=(min(tnow)+max(tnow))/2;
            %population must be within 0.01 Aitchison distance from the
            %pre-kick state (this will be too small if we study a larger
            %population)
        end

        if tn(i)>tu_n/4
            if (max(max([log(XVmax_n./XVmax);log(XVmin./XVmin_n)],[],1).*...
                    (XVeq_n>10^-4*sum(XVeq_n)).*(XVstd_n./XVeq_n>0.01))<th2 &...
                    ret_time(2)==0)
                ret_time(2)=(min(tnow)+max(tnow))/2;
                %they have to be within the bounds of the previous
                %system, or the species must be rare (<10^-4 proportion) or
                %stationary (CoV<0.01)
            end
            if (max(log(XVstd_n./XVstd).*(XVeq_n>10^-4*sum(XVeq_n)).*(XVstd_n./XVeq_n>0.01))<th2 &...
                    ret_time(3)==0)
                %if they are fluctuating they must have amplitudes less
                %than th2 of their previous final, OR the fluctations must
                %be less than 0.01 of the population size, OR the
                %population size must be less than 10^-4 of the total
                %population
                ret_time(3)=(min(tnow)+max(tnow))/2;
            end
            if ((XVwcov_n<0.001 | XVwcov_n<1.01*tot_victim_wcov) &...
                    ret_time(5)==0)
                %the weighted CoV must be within 1% of previous final, or
                %less than 0.001
                ret_time(5)=(min(tnow)+max(tnow))/2;
            end
        end
        i=i+1;
        if tn(i)+tu_n/2>size(soln,1)-1
            ret_time(ret_time==0)=Inf;
            break
        end
    end
    %     end

    rt_vect=[rt_vect;ret_time];

end
```
