## Supplementary material for "Lysogeny destabilizes computationally simulated microbiomes": a.Codes and data for Gilman et al 2023.zip: readme.txt

This folder contains the following files:

Simulation codes:

microbiome_simulator.m (MATLAB): This code simulates microbiome dynamics using the model described in Gilman et al., measures return times to the mature state of each microbiome after simulated disturbances, and records the results. Data reported in the paper correspond to simulations with seed numbers 1 to 20000 for each parameter combination studied. This code calls functions get_dVP_nm.m (constructs dynamical systems), v_and_p_func_nm.m (runs simulations to maturity),  hello_chaos_nm.m (tests whether the dynamics of mature states are chaotic), and qcc_func_nm.m (measures return times after disturbances).

Data:

spec_no_dd_full.csv (text): simulation results when phages are specialists and lysogeny and induction are not density dependent, as reported in the body of the paper

gen_no_dd_full.csv (text): simulation results when phages can be generalists and lysogeny and induction are not density dependent, as reported in the supplementary information

spec_dd_full.csv (text): simulation results when phages are specialists and lysogeny and induction can be density dependent, as reported in the supplementary information

Analysis codes:

make_surfs.m (MATLAB): uses the data provided to reproduce figures 1-3, S2, and S3

analyse_results_specialists.R (R): conducts regression models to reproduce results in Table 1 and Table S4, and reproduces figure S5 

analyse_results_generalists.R (R): conducts regression models to reproduce results in Table S2 and Table S4, and reproduces figure S4 

analyse_results_dd.R (R): conducts regression models to reproduce results in Table S3 and Table S4, and reproduces figure S6 
