## Supplementary material for "Lysogeny destabilizes computationally simulated microbiomes": a.Codes and data for Gilman et al 2023.zip: v_and_p_func_nm.m

```
function [Xfinal,Xeq,XVfinal,XVeq,XVmin,XVmax,XVstd,XVclr,tot_victim_sr,tot_victim_shan,tot_victim_wcov,Alpha,K,A,mp,tu,lys_apt,dVP,mean_lys]=v_and_p_func_nm(Pv,Pp,nv,cv,Kmax,r,mh,w,np,att,z,cp,b,d,f1,f2,g1,x,g2,g3,c,s,lp)

%Create the competiton matrix
V=randn(Pv,nv);
disttri=pdist(V);
Atri=exp(-disttri.^2/cv);
Alpha=squareform(Atri)+eye(Pv);
%Make competition matrix asymmetircal 
Af=exp((ones(size(Alpha))-eye(size(Alpha,1))).*randn(size(Alpha))*0.01);
Alpha=Alpha.*Af;

%Set carrying capacities
K=Kmax*exp(-(V.^2)*ones(nv,1));

%Control which viruses enter system by setting some mirgation rates to zero
mp=(mh/10)*randsample([ones(Pp,1);zeros(Pv-Pp,1)],Pv);   %migration rate for predators

%Create the attack rate matrices for virulent and temperate viruses
Ppos=V(:,1:np);     %position of predators in trait space
disttriPV1=squareform(pdist([V(:,1:np);Ppos]));
disttriPV=disttriPV1(1:Pv,1+Pv:2*Pv);
pred_apt=exp(-disttriPV.^2/cp); %the ecological suitability of each prey species
pred_sten=sign((pred_apt==1)+(rand(size(pred_apt))<z));
pred_sten=pred_sten.*(ones(Pv,1)*(mp>0)');
A=att*pred_apt.*pred_sten;
lys_apt=(mp>0).*(rand(Pv,1)<lp);

%Now, we have everything we need to compute the vector of population growth
%rates. In order to solve this with MATLAB's ODE solvers, we will need to
%write the growth rates as a function of the parameters and the vector of
%species densities X. The funciton is complicated, so I will create it
%as a separate script.
options_coarse=odeset('NonNegative',[1:(3*Pv)]);
options_fine=odeset('NonNegative',[1:(3*Pv)],'RelTol',1e-6,'AbsTol',1e-8);
dVP=@(t,X) get_dVP_nm(X,Alpha,K,r,mh,mp,A,b,d,w,f1,f2,g1,x,g2,g3,c,s,lys_apt);

%Now, let's solve this starting from an empty system, with a bump at
%t=1000. The goal here is to make sure that if there is some sort of
%idiosyncratic state that can be reached by starting with an empty system,
%we get bumped away from that state. We never found such an idiosyncratic
%state, but if we take out calls to the random number generator now, our
%results will not repeat exactly.
X=zeros(3*Pv,1);
[t,sol] = ode15s(dVP,[0 1000],X,options_coarse);
X=(sol(size(sol,1),:).*exp(0.1*randn(1,3*Pv)))';

tu=200; %set the number of timesteps over which to consider the final state
[t,sol] = ode15s(dVP,[0:5000],X,options_fine);
Xfinal=sol(t>=t(end)-tu,:);
Xeq=mean(Xfinal,1);
XVfinal=Xfinal(:,1:Pv)+Xfinal(:,1+2*Pv:3*Pv);

%compute mean for hosts an lysogens
XVeq=mean(XVfinal,1);

%compute mean lysogeny weighted by step length 
lys_probs=(1-(1-f1)*exp(-(2*Xfinal(:,1+Pv:2*Pv)))).*(ones(size(Xfinal,1),1)*lys_apt');
mean_lys=sum(sum(lys_probs.*Xfinal(:,1:Pv).*Xfinal(:,1+Pv:2*Pv)))/...
    sum(sum(Xfinal(:,1:Pv).*Xfinal(:,1+Pv:2*Pv).*(ones(size(Xfinal,1),1)*lys_apt')));

XVmax=max(XVfinal,[],1);
XVmin=min(XVfinal,[],1);
XVstd=std(XVfinal,[],1);
XVclr=log(XVeq)-mean(log(XVeq));

tot_Vprop=(Xeq(1:Pv)+Xeq(1+2*Pv:3*Pv))/sum(Xeq(1:Pv)+Xeq(1+2*Pv:3*Pv));
tot_victim_sr=sum(tot_Vprop>10^-4);
tot_victim_shan=-sum(tot_Vprop.*log(tot_Vprop));
tot_victim_wcov=sum(XVstd)/sum(mean(XVfinal));
```
